# A 6-nm ultra-photostable DNA Fluorocube for fluorescence imaging

**DOI:** 10.1101/716787

**Authors:** Stefan Niekamp, Nico Stuurman, Ronald D. Vale

## Abstract

Photobleaching limits extended imaging of fluorescent biological samples. Here, we developed DNA origami-based “Fluorocubes” that are similar in size to the green fluorescent protein (GFP), have single-point attachment to proteins, have a 50-fold higher photobleaching lifetime and emit 40-fold more photons than single organic dyes. We demonstrate that DNA Fluorocubes provide outstanding tools for single-molecule imaging, allowing the tracking of single motor proteins for >800 steps with nanometer precision.

## Introduction

Imaging proteins and macromolecular complexes at the single-molecule level is a powerful method to study distribution, stoichiometry, dynamics and precise motion of molecular machines^1^. To achieve high spatiotemporal resolution, proteins of interest are often labeled with luminescent probes^1^. An ideal probe is photostable, (continuously) bright, small, and can be monovalently attached to biological molecules. While organic dyes^2,3^ and fluorescent proteins^4^ fulfill the latter two criteria, they often suffer from photobleaching, which leads to a low signal. Alternative probes such as quantum dots^2,5^ and other fluorescent nanoparticles^6^ are very bright and extremely photostable, but frequently exhibit large fluctuations in intensity (blinking). The relatively large (∼15 nm) size of these probes (**Supplementary Fig. 1 a**) can perturb protein function and complicates multicolor labeling of proteins^2,3,5^. Large probes often lack control over surface chemistry, which can lead to multiple proteins attaching to the same fluorescent probe^7^. Here, we developed small (6 nm) DNA-based Fluorocubes that have single-point attachment, exhibit continuous emission, emit up to ∼40-fold more photons than single organic dyes, and are ∼50-fold more photostable than single organic dyes.

## Results

Previous work established that organic dyes placed within 2 nm result in quenching^7^, while dye spacing of >5 nm results in a linear intensity increase with the number of dyes^8^. We were interested in the properties of probes separated by 2 to 5 nm.

To set the position of dyes with nanometer precision, we took advantage of tools from DNA nanotechnology^8–10^. Using the scaffold-free DNA origami approach^11^, we designed a DNA Fluorocube composed of four 16 base-pair (bp) long double-stranded DNA helices with organic dyes positioned at six of its eight corners (**Fig. 1 a, b, Supplementary Fig. 1 b**). We reserved one position for the placement of a HALO-ligand, benzylguanine (for SNAP tag), thiol, biotin, or amine to label proteins at specific locations (**Fig. 1 b**).

**Figure 1.**
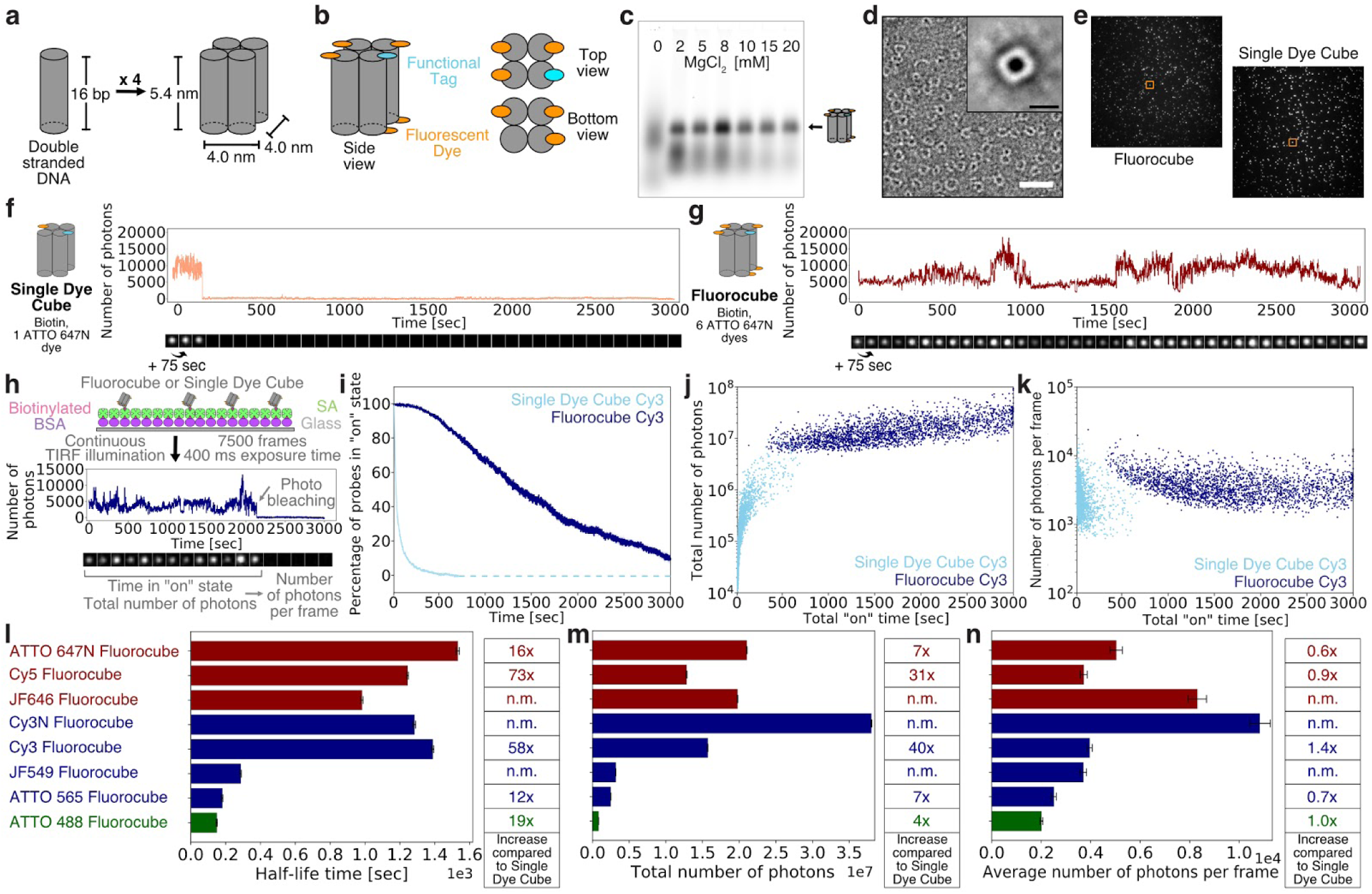
|Design, assembly, and photophysical properties of DNA Fluorocubes. (**a**) Design and shape of DNA Fluorocubes. Four 32 bp long single-stranded DNAs (ssDNA) are connected using crossovers resulting in four connected 16 bp long double-stranded DNAs (dsDNA) with a size of approximately 5.4 × 4.0 × 4.0 nm. A detailed map of oligonucleotide routing and the bases is depicted in **Supplementary Figure 1 b**. (**b**) Each of the 5’ and 3’ ends of the DNA can be functionalized. For the DNA Fluorocube design we used six fluorophores and one functional tag such as HALO-ligand, benzylguanine for SNAP tag, thiol, biotin, or amine. (**c**) 2% agarose gel of DNA Fluorocubes after thermal annealing. The four ssDNA strands are annealed with different MgCl_2_ concentrations. Quantification of assembly yield is shown in **Supplementary Figure 1 c**. Here the ssDNAs were modified with six ATTO 647N dyes and one biotin. (**d**) Negative stain electron microscopy image of DNA Fluorocubes. Insert shows class average of 983 particles. More images are shown in **Supplementary Figure 1 d and Supplementary Figure 2**. Here the Fluorocube was labeled with six ATTO 647N dyes and one biotin. White scale bar: 30 nm. Black scale bar: 6 nm. (**e**) TIRF image of a biotin functionalized ATTO 647N Fluorocube (Fluorocube) and a cube with one ATTO 647N dye (Single Dye Cube). Orange boxes show molecules whose intensity traces are shown in **f** (Single Dye Cube) and **g** (Fluorocube). (**f**) Example intensity trace of ATTO 647N Single Dye Cube with one biotin. (**g**) Example intensity trace of ATTO 647N Fluorocube with one biotin. (**f, g**) More intensity traces with different fluorophores are shown in **Supplementary Figure 3**. (**h**) Experimental setup for quantification of photophysical properties of Fluorocubes and Single Dye Cubes. Biotinylated Fluorocubes and Single Dye Cubes are attached to a cover slip (Glass) with biotinylated BSA, and streptavidin (SA). Then movies with 7,500 frames total and 400 ms exposure each are recorded under continuous laser illumination. Intensity traces of single-molecules are analysed for time in “on” state, total number of photons and number of photons per frame. A detailed description of the analysis can be found in **Online Methods**. (**i**) Photostability of Cy3 Fluorocubes and Cy3 Single Dye Cubes. The survival rate was quantified by counting the percentage of probes in the “on” state at any given time from 0 to 3,000 seconds. Opaque color is the standard error of the mean of four repeats with more than 500 molecules each. Once all probes photobleached data analysis was terminated. This is indicated by the dashed line. (**j**) Total number of photons of Cy3 Fluorocubes and Cy3 Single Dye Cubes as a function of the total “on” time at the single-molecule level. (>**k**) Average number of photons per frame of Cy3 Fluorocubes and Cy3 Single Dye Cubes as a function of the total “on” time at the single-molecule level. (**l**) Histogram of the half-life time of DNA Fluorocubes with different fluorophores. (**m**) Histogram of the total number of photons of DNA Fluorocubes with different fluorophores. Note, that the average of the total number of photons will be slightly higher than shown here because not all probes bleached within 3,000 seconds. (**n**) Histogram of the average number of photons per frame of DNA Fluorocubes with different fluorophores. (**l-n**) The tables on the right show fold increase of DNA Fluorocubes compared to Single Dye Cubes. The error bars show the standard error of the mean of four repeats with more than 500 molecules each. Single-molecule distributions are shown in **Supplementary Figure 5, 6, and 11**. “Cy3” stands for the non-sulfonated version of Cy3 whereas “Cy3N” stands for the sulfonated version of Cy3. “n.m.” is not measured.

Since small DNA nanostructures are difficult to assemble because of high electrostatic repulsion between the negatively-charged DNA strands^12^, we optimized the folding yield and determined the structural integrity of the DNA Fluorocubes after thermal annealing. Using agarose gel electrophoresis, we measured a folding yield of over 60% (**Fig. 1 c, Supplementary Fig. 1 c**). Negative stain transmission electron microscopy (TEM) showed that Fluorocubes assembled into the desired shape (**Fig. 1 d, Supplementary Fig. 1 d**). However, we noticed some dye-dependent shape changes; Fluorocubes with ATTO dyes assembled into cubes with the predicted size, whereas Fluorocubes made with non-sulfonated cyanine dyes folded into slightly larger cubes with a hollow center (**Supplementary Fig. 2**). Overall, the protocol for the assembly of DNA Fluorocubes is easy and optimized for high yield.

Next, we compared DNA Fluorocubes to one single dye attached to the same DNA-origami scaffold (Single Dye Cube) for several different dyes such as ATTO 488, ATTO 565, ATTO 647N, Cy3, or Cy5. Surface-immobilized Fluorocubes and Single Dye Cubes were imaged using total internal reflection fluorescence (TIRF) microscopy (**Fig. 1 e-h**). Single Dye Cubes showed roughly constant intensity over time and then one-step photobleaching disappearance (**Fig. 1 f, Supplementary Fig. 3**). Fluorocubes, on the other hand, often increased in brightness during the time of acquisition (**Fig. 1 g, Supplementary Fig. 3**).

Using the intensity traces of many Fluorocubes and Single Dye Cubes, we quantified the total “on” time as a measure of photostability, the total number of photons emitted, the average number of photons per frame as a measure of brightness, and the coefficient of variance as a measure of blinking (**Fig. 1 h, Supplementary Fig. 4 a**). Overall, Fluorocubes are significantly more photostable than Single Dye Cubes. For instance, after 99% of the Cy3 Single Dye Cubes bleached, more than 80% of the Cy3 Fluorocubes were still in the “on” state, resulting in an increased half-life of more than 50-fold (**Fig. 1 i, l**). However, we noticed a strong dye-to-dye variability (**Fig. 1 l, Supplementary Fig. 5, Supplementary Movies 1-5, Supplementary Table 3**). While Cy3 Fluorocubes had about 90% of probes in the “on” state after 600 sec of continuous 561 nm laser illumination (120 W/cm^2^), ATTO 565 Fluorocubes had only about 10% of probes in the “on” state (**Fig. 1 i, Supplementary Fig. 5 a**). As expected based upon this increased photostability, the total number of photons per Fluorocube before photobleaching was up to 40-fold higher compared to Single Dye Cubes (**Fig. 1 j, m**). specifically, Cy3 Fluorocubes emitted an average of 1.57 ± 0.04 × 10^7^ photons, whereas Cy3 Single Dye Cubes emitted 3.90 ± 0.10 × 10^5^ photons, similar to previous reports^13^.

Even though there are 6-fold more dyes, the brightness of the Fluorocubes was similar to the Single Dye Cubes (**Fig. 1 k, n, Supplementary Fig. 5**). The Fluorocubes displayed a slight increase in the coefficient of variance of photon emission (**Supplementary Fig. 4**). This slight increase is not caused by dark states like for quantum dots^2,7^, but rather by brightness increases during imaging (**Supplementary Fig. 3**). In addition to the dyes listed above, we also tested Fluorocubes with Janelia Fluorophores JF549 and JF646^14^ and found that JF646 Fluorocube has a similar increase in photostability to the Cy3 Fluorocube, while the JF549 Fluorocube shows a much smaller gain in photostability (**Fig. 1 l-n, Supplementary Fig. 6**). Taken together, most DNA Fluorocubes have significantly greater photostability than single dyes (**Fig. 1 l-n**) and emit up to 40-fold more photons.

Since quantum dots are very bright and photostable, we compared the photophysical properties of ATTO 647N Fluorocubes to 655 Qdot Nanocrystals. While these quantum dots emit about four-fold more photons and are about four-fold brighter on average than ATTO 647N Fluorocubes, quantum dots blink significantly more than DNA Fluorocubes (**Supplementary Fig. 7**). Moreover, quantum dots tend to enter dark states in which they do not emit photons for a couple of seconds (**Supplementary Fig. 7 a**), making it difficult to track quantum dots continuously.

Next, we examined how DNA Fluorocubes respond to a wide range of excitation powers and whether they are influenced by oxygen scavenger^15^ or triplet quencher^16^. Fluorocubes showed a linear increase in intensity until saturation, similar to Single Dye Cubes (**Supplementary Fig. 8 a-c**) and performed better with oxygen scavenger and triplet quencher (**Supplementary Fig. 8 d-f**). However, even without these aids, Fluorocubes outperformed Single Dye Cubes with the oxygen scavenger system and triplet quencher. We also measured absorption and emission spectra (**Supplementary Fig. 9**), fluorescence lifetime and anisotropy (**Supplementary Fig. 10**), but we did not find any parameter that correlated with a change in photostability of dyes when coupled to a Fluorocube.

The hole sizes in different DNA Fluorocubes, as seen with negative stain TEM (**Supplementary Fig. 2**), suggested charge may influence the assembly and performance of Fluorocubes. We therefore compared sulfonated Cy3 dyes to the non-sulfonated version described earlier. By TEM, we saw that the hole in the DNA origami was smaller for the sulfonated Cy3 (**Supplementary Fig. 2, Supplementary Table 3**). The sulfonated Cy3 emitted twice as many photons as the non-sulfonated Cy3 while having similar photostability (**Fig. 1 l, m, Supplementary Fig. 11, Supplementary Table 3**). These data show that charge and hydrophobicity of cyanine dyes attached to the DNA Fluorocube scaffold influence its performance.

**Figure 2.**
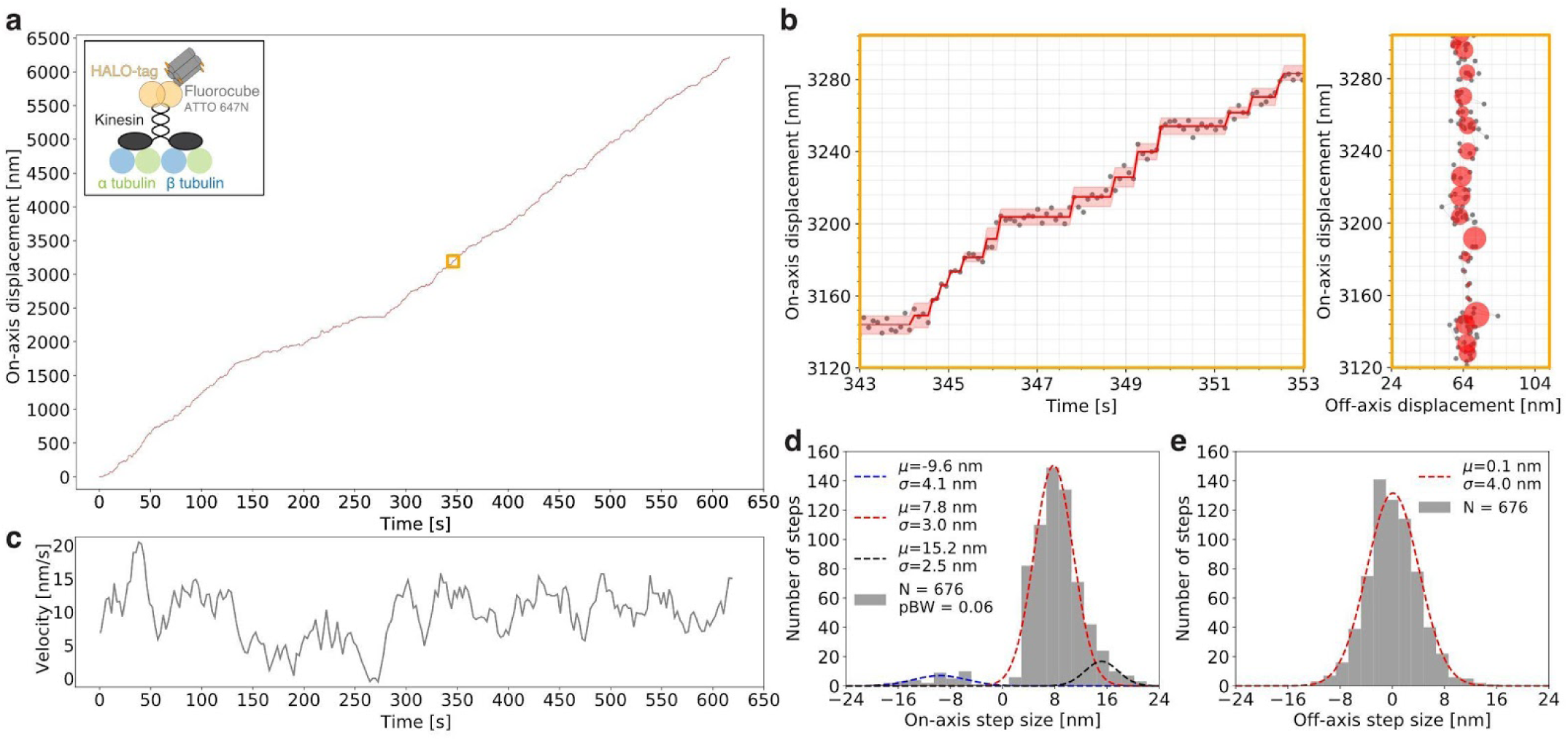
Tracking steps of a single kinesin over more than 6 µm. (**a**) Raw stepping data with position versus time of a single kinesin (grey dots) over 6 µm with detected steps (red line) along an axoneme. The opaque red line shows the standard deviation for each step. The insert shows the experimental setup for which a kinesin is labeled with one ATTO 647N Fluorocube with a HALO-tag (for details see **Online Methods**). The orange box is enlarged in **b**. The movement of this particular kinesin is shown in **Supplementary Movie 6**. (**b**) Left: Zoom-in on the raw stepping data with position versus time of a single kinesin as shown in **a**. Right: Same trace as on the left but in XY space. The grey dots are raw data and the red circles show the fitted position for which the radius corresponds to the standard deviation. (**c**) Velocity over time for the stepping trace of a single kinesin as shown in **a**. The grey line shows a moving average of velocity binned into 15.6 sec (for details see **Online Methods**). (**d**) Histogram of the on-axis step size distribution of the data shown in **a**. The data was split into positive and negative steps and fit with Gaussians. For the negative steps, a single Gaussian was used (blue) whereas for the positive steps two Gaussians were used (red, black). pBW is the fraction of backward steps. (**e**) Histogram of the off-axis step size distribution of the data shown in **a** fitted with a single Gaussian. (**d, e**) We detected 821 steps but only used 676 steps for further quantification because we only counted steps for which the step itself and the previous as well as following step had a standard deviation of less than 4 nm. Additional stepping traces and quantification are shown in **Supplementary Figure 12**.

We next tested whether Fluorocubes can be attached to proteins and used for prolonged readouts of activity. For this purpose, we labeled an ultra processive kinesin KIF1A^17^ with ATTO 647N Fluorocubes using a C-terminal HALO-tag (**Fig. 2 a**) and imaged it moving along axonemes. Labeling kinesin with a single Fluorocube enabled us to record more than 800 steps of an individual motor with nanometer precision (**Fig. 2 a, b, Supplementary Movie 6**). The trace in Figure 2 revealed an on-axis step size of 7.8 nm with almost no off-axis stepping (**Fig. 2 d, e**) which is in good agreement with previous reports^17,18^ (discussion in **Supplementary Fig. 12**). Thus, Fluorocubes do not interfere with protein function. Moreover, using Fluorocubes allowed us to collect more than 6,000 data points of an individual motor compared to approximately 200 data points that can be collected at a similar resolution with a single organic dye^19,20^. By recording very long traces, we could detect occasional pausing and velocity fluctuations within the trace of an individual kinesin (**Fig. 2 c**). Previous work demonstrated that different kinesins can have different velocities^21^, but our prolonged observations showed that even individual kinesins undergo considerable velocity fluctuations over time.

## Discussion

In summary, we developed small (6-nm), ultra-photostable fluorescent probes. We have shown that some dyes - bound to DNA Fluorocubes - emit over 40-fold more photons than single organic dyes. Further improvement of DNA Fluorocubes could be made by using other dyes such as self-healing fluorophores^13^, by adding DNA intercalating dyes^22^, by increasing the number of fluorophores on the Fluorocube^8^, by changing the spacing between dyes (either by changing the cube size or by alternating the dye linker length)^7,12^, or by changing the DNA sequence close to the fluorophores^23^. A full understanding of the photo-physical mechanisms underlying DNA Fluorocube photostability remains to be uncovered but we speculate that resonance energy transfer between individual dyes in a Fluorocube plays an important role in this phenomenon.

DNA Fluorocubes are easily prepared from commercially available reagents and can be attached to all commonly used protein tags, making them simple to use for in vitro studies. Here, we measured the walking of kinesin for extended periods of time, without any evidence of perturbation by the probe. We anticipate that DNA Fluorocubes will become the reagent of choice for extracellular single-molecule imaging experiments. Beyond single-molecule studies, the long photobleaching-lifetime and high number of total photons of DNA Fluorocubes could prove useful in numerous other fluorescence imaging applications, including FISH^24^, MERFISH^25^, DNA-PAINT^26^, or immunofluorescence for research and medical diagnosis.

## Methods

### Flow-cell preparation

Flow-cells were assembled as previously described^27^. Briefly, we made custom three-cell flow chambers using laser-cut double-sided adhesive sheets (Soles2dance, 9474-08×12 - 3M 9474LE 300LSE), glass slides (Thermo Fisher Scientific, 12-550-123), and 170 μm thick coverslips (Zeiss, 474030-9000-000). The coverslips were cleaned in a 5% v/v solution of Hellmanex III (Sigma, Z805939-1EA) at 50° C overnight and washed extensively with Milli-Q water afterwards.

### Assembly and analysis of DNA Fluorocubes

For each Fluorocube four 32 bp long oligonucleotide strands are required, each modified either with dyes or a functional tag such as biotin or HALO-ligand^28^ (**Supplementary Fig. 1b, Supplementary Table 1, 2**). These four oligos were mixed to a final concentration of 40 μM in folding buffer (5 mM Tris pH 8.5, 1 mM EDTA and 40 mM MgCl_2_) and annealed by denaturation at 85° C for 5 min followed by cooling from 80° C to 65° C with a decrease of 1° C per 5 min. Afterwards the samples were further cooled from 65° C to 25° C with a decrease of 1° C per 20 min and then held at 4° C. Folding products were analyzed by 2.0% agarose gel electrophoresis in TBE (45 mM Tris-borate and 1 mM EDTA) with 12 mM MgCl_2_ and purified by extraction and centrifugation in Freeze ‘N Squeeze columns.

Agarose gel-based yield estimation was carried out using <u>ImageJ</u>^29^. The percentage of Fluorocubes that ran as a monomeric band was estimated as the background-subtracted integrated intensity value divided by the background-subtracted integrated intensity value enclosing the material from the well, down to the bottom of the leading band (single oligos).

### Negative stain electron microscopy data collection and processing

For negative-stain EM, unpurified, but folded Fluorocubes at 300 nM were incubated on freshly glow discharged carbon coated 400 mesh copper grids for 1 min and blotted off. Immediately after blotting, a 0.75% uranyl formate solution was applied for staining and blotted off without incubation. This staining was repeated four times and followed by a last incubation for which the stain was incubated for 45 sec before blotting. Samples were allowed to air dry before imaging. Data were acquired at UCSF, on a Tecnai T12 microscope operating at 120 kV, using a 4k×4k CCD camera (UltraScan 4000, Gatan) and a pixel size of 1.7 Å/pixel. For the class average in **Figure 1 d**, 1,743 Particles were picked and boxed using EMAN 2.21^30^. Then a 2D classification was performed to remove junk and noisy particles, leading to 983 particles selected.

### Mono-Q clean-up of DNA Fluorocubes

Thermally annealed DNA Fluorocubes were purified using anion exchange chromatography with a GE Source 15Q 4.6/100 PE column (**Supplementary Fig. 13**). DNA Fluorocubes were bound to the column in Buffer A (20 mM Tris pH 8.0, 1 mM EDTA, 10 mM Mg-Ac, and 10% Glycerol) and afterwards the ionic strength was increased linearly by adding Buffer B (20 mM Tris pH 8.0, 2 M K-Ac, 1 mM EDTA, 10 mM Mg-Ac, and 10% Glycerol) to 100% over 80 min.

### Labeling of oligonucleotides with Janelia Fluorophores

We mixed 5’ and 3’ amino modified oligos (**Supplementary Table 1**) at a final concentration of 500 µM (in water) with NHS ester modified Janelia Fluorophores JF549 or JF646^14^ at a final concentration of 5 mM (in DMSO) in 15 mM HEPES pH 8.5 buffer. These solutions were incubated for 4 h at room temperature. We then removed excess dye by four subsequent spins of the solution over Micro Bio-Spin 6 Columns (Bio-Rad) at 700 g for 2 min. The final oligo concentration was determined with a UV spectrophotometer. Afterwards Fluorocubes were assembled as described above.

### Preparation of flow-cells with DNA Fluorocubes, Single Dye Cubes and quantum dots

The preparation of flow cells is identical for 6-dye cubes (Fluorocubes) or 1-dye Cubes (Single Dye Cubes). In either case, the Fluorocubes were folded with biotin as the functional tag. We first added 10 µl of 5 mg/ml Biotin-BSA in BRB80 to the flow-cell and incubated for 2 min. Afterwards, we added another 10 µl of 5 mg/ml Biotin-BSA in BRB80 and incubated for 2 min. Then we washed with 20 µl of Fluorocube Buffer (20 mM Tris pH 8.0, 1 mM EDTA, 20 mM Mg-Ac, and 50 mM NaCl) with 2 mg/ml -casein (Sigma, C6905), 0.4 mg/ml κ-casein (Sigma, C0406). This was followed by addition of 10 µl of 0.5 mg/ml Streptavidin in PBS (pH 7.4) and a 2 min incubation. We then washed with 20 µl of Fluorocube Buffer with 2 mg/ml -casein, and 0.4 mg/ml κ-casein. Next, we either added Fluorocubes in Fluorocube Buffer or Quantum dots (Qdot™ 655 Streptavidin Conjugate, ThermoFisher Scientific, Q10121MP) and incubated for 5 min. Finally, we washed with 30 µl of Fluorocube Buffer with 2 mg/ml -casein, and 0.4 mg/ml κ-casein. We then added the PCA/PCD/Trolox oxygen scavenging system^15,16^ in Fluorocube Buffer with 2 mg/ml -casein, and 0.4 mg/ml κ-casein for the Fluorocubes. For the Quantum dots we added the PCA/PCD oxygen scavenging system^15^ and 1% β-mercaptoethanol in Fluorocube Buffer with 2 mg/ml -casein, and 0.4 mg/ml κ-casein.

### Measurements of anisotropy, lifetime, absorption and emission spectra

We determined anisotropy, lifetime, as well as absorption and emission spectra using an ISS K2 multifrequency fluorometer in bulk measurements. All experiments were performed at room temperature (21-23° C). Instrument settings for the anisotropy measurements are listed in **Supplementary Table 5**, settings for the fluorescence lifetime measurements are listed in **Supplementary Table 6**, and settings for the absorption and emission spectra are listed in **Supplementary Table 7**.

### Kinesin cloning, purification and labeling

The kinesin construct was cloned and purified as previously described^17^ except that the GFP was replaced with a HALO-tag^28^.

The plasmid was transfected and expressed in Agilent BL21(DE3) cells. Cells were grown in LB at 37° C until they reached 1.0 OD_280_ and the expression was induced by addition of 0.2 mM IPTG. Then the cells were incubated overnight at 20° C. Cells were pelleted and harvested in lysis buffer (25 mM Pipes (pH 6.8), 2 mM MgCl_2_, 250 mM NaCl, 20 mM imidazole, 1 mM BME, 0.1 mM ATP, and 0.4 mM PMSF), and lysed in the EmulsiFlex homogenizer (Avestin). After a spin in a Sorvall SS-34 rotor for 30 min at 30,000 × g, the supernatant was loaded onto a Ni-NTA resin (QIAGEN, 30210) and washed with additional lysis buffer. We then took 500 μl of beads slur in lysis buffer supplemented with 10 mM MgCl_2_ and added ATTO 647N Fluorocubes with a HALO-tag^28^ to 5 μM final. This mixture was incubated on a Nutator for 3 h at 4° C. Afterwards we washed with additional lysis buffer supplemented with 10 mM MgCl_2_. Then the protein was eluted by adding 300 mM of imidazole to the lysis buffer supplemented with 10 mM MgCl_2_. Subsequently the sample was purified by gel filtration over a S200 10/300GL column from GE Healthcare. Gel filtration buffer was composed of 25 mM Pipes (pH 6.8), 10 mM MgCl_2_, 200 mM NaCl, 1 mM EGTA, 1 mM DTT, and 10% sucrose. Finally the sample was flash frozen and stored at −80° C.

### Preparation of flow-cells with kinesin

Single-molecule assays with kinesin in flow-cells were prepared as previously described^17,19^. We first added 10 µl of Alexa 488 labeled axonemes in BRB80 (80 mM Pipes (pH 6.8), 1 mM MgCl_2_, 1 mM EGTA) and incubated for 5 min. Then, we washed with 60 µl of BRB80 with 1.0 mg/ml κ-casein (Sigma, C0406) supplemented with 5 mM MgCl_2_. Next, we added 10 µl of ATTO 647N Fluorocubes labeled kinesin in BRB80 with additional 5 mM MgCl_2_, 1.0 mg/ml κ-casein, 1.5 µM ATP, an ATP regeneration system (1 mM phosphoenolpyruvate (Sigma), 0.01 U pyruvate kinase (Sigma), 0.03 U lactate dehydrogenase (Sigma)), and the PCA/PCD/Trolox oxygen scavenging system^15,16^.

### Microscope setup

All data collections were carried out at room temperature (∼23° C) using a total internal reflection fluorescence (TIRF) inverted microscope (Nikon Eclipse Ti microscope) equipped with a 100× (1.45 NA) oil objective (Nikon, Plan Apo ƛ). We used an Andor iXon 512×512 pixel EM camera, DU-897E and a pixel size of 159 nm. We used two stepping motor actuators (Sigma Koki, SGSP-25ACTR-B0) mounted on a KS stage (KS, Model KS-N) and a custom-built cover to reduce noise from air and temperature fluctuations. A reflection based autofocus unit (FocusStat4) was custom adapted to our microscope (Focal Point Inc.). For the data collection we used a 488 nm laser (Coherent Sapphire 488 LP, 150 mW), a 561 nm laser (Coherent Sapphire 561 LP, 150 mW), and a 640 nm laser (Coherent CUBE 640-100C, 100 mW). A TIRF cube containing excitation filter (Chroma, zet405/491/561/638x), dichroic mirror (zt405/488/561/638rpc), and emission filter (Chroma, zet405/491/561/647m) was mounted in the upper turret of the microscope. The lower turret contained an ET450/50m (Chroma) filter for the 488 nm laser, an ET600/50m (Chroma) filter for the 561 nm laser, and an ET700/75m (Chroma) filter for the 640 nm laser.

### Single-molecule TIRF data collection and analysis of DNA Fluorocubes and Single Dye Cubes

The TIRF data of surface immobilized DNA Fluorocubes and Single Dye Cubes was acquired under continuous laser illumination with an intensity (irradiance) of 120 W/cm^2^ (488 nm laser), 120 W/cm^2^ (561 nm laser), and 160 W/cm^2^ (640 nm laser) and an exposure of 400 ms if not specified otherwise. The acquisition length varied based on the experiment. Typically we either recoded 7,500 frames (3,000 sec), 1,500 frames (600 sec), or 300 frames (120 sec). We used the camera in conventional CCD mode (i.e., no EM gain). All datasets were acquired with a ‘16 bit, conventional, 3 MHz’ setting and a preamp gain of 5x. All experiments were performed at room temperature (21-23° C). The acquisition software was μManager^31^ 2.0 and data was analyzed in ImageJ^29^. Single molecules were located and traced using the <u>Spot Intensity Analysis</u> plugin in ImageJ^29^ with the following settings: Time interval of 0.4 sec (except for data in **Supplementary Figure 8** for which the exposure time was used as listed in **Supplementary Table 8**), Electron per ADU of 1.84, Spot radius of 3, Noise tolerance of 100, and a Gaussian background estimation. The number of frames to check is shown in **Supplementary Table 9** since it varies for each sample depending on how fast they bleach. Afterwards the data was analyzed with a custom written python script.

### Single-molecule TIRF data collection and analysis of kinesin stepping

Kinesins labeled with ATTO 647N Fluorocubes with a HALO-tag^28^ were continuously illuminated with a 640 nm laser (160 W/cm^2^) with an effective exposure time of 0.103 s. We used the camera in conventional CCD mode (i.e., no EM gain). All datasets were acquired with a ‘16 bit, conventional, 3 MHz’ setting and a preamp gain of 5x. All experiments were performed at room temperature (21-23° C). The acquisition software was μManager^31^ 2.0. All emitters were fitted and localized using μManager’s^31^ “Localization Microscopy’ plug-in as previously described^27^. Parameters for analysis are shown in **Supplementary Table 10**. Tracks of individual motors were extracted using the μManagers^31^ “Localization Microscopy’ plug-in. We set the minimum frame number to 1000, the maximum number of missing frames to 100, the maximum distance between frames to 100 nm and the total minimum distances of the full track to 500 nm. Then tracks of individual motors were rotated using a principal component analysis (PCA) implemented in python. Afterwards we used a custom Matlab (Matlab R2016b) script to identify individual steps using Chung-Kennedy edge-detecting algorithm as previously described^32^ and further analyzed the data in a custom written python script. Only steps for which the step itself and the previous as well as following step had a standard deviation of less than 4 nm were considered for further quantification. The velocity over time for the stepping trace of a single kinesin was analyzed with a moving average for which we first binned the data into 2.6 sec bins and then grouped six of these 2.6 sec bins into 15.6 sec bins. We used these 15.6 sec bins to calculate the average velocity at any given point, but used the 2.6 sec intervals to move along the time axis.

### Figure preparation

Figures and graphs were created using ImageJ^29^ (light microscopy data), Affinity designer (version 1.6.1, Serif (Europe) Ltd) and Python (version 2.7, Python Software Foundation).

### Statistics

For each result the inherent uncertainty due to random or systematic errors and their validation are discussed in the relevant sections of the manuscript. Details about the sample size, number of independent calculations, and the determination of error bars in plots are included in the figures and figure captions.

## Code availability

μManager acquisition and analysis software is available partly under the Berkeley Software Distribution (BSD) license, partly under the GNU Lesser General Public License (LGPL) and development is hosted on GitHub at https://github.com/nicost/micro-manager. The latest version for MacOS and Windows can be downloaded here: https://valelab.ucsf.edu/~nico/mm2gamma/. All other code is available from the authors upon request.

## Data availability

Example raw datasets of the DNA Fluorocubes used to determine the photophysical properties are hosted on Zenodo: https://doi.org/10.5281/zenodo.3350875. All other data files are available from the authors upon request.

## Acknowledgements

We are grateful to Jongmin Sung for critical discussions of the manuscript. We thank Dyche Mullins for teaching us how to use the ISS K2 multifrequency fluorometer. We thank Luke Lavis (Janelia Research Campus) for the suggestion of comparing sulfonated versus non-sulfonated dyes and for providing the JF549 and JF646 dyes. Andrew Carter and Elizabeth Villa supplied the Matlab script for step detection of kinesin. The authors gratefully acknowledge funding from the National Institutes of Health: R01GM097312, 1R35GM118106 (R.D.V., S.N.), and the Howard Hughes Medical Institute (R.D.V. and N.S.).

## Author contributions

S.N., N.S., and R.D.V. designed the research; S.N. prepared samples; S.N. collected TIRF microscopy data; S.N. collected electron microscopy data; S.N. analyzed the data; S.N., N.S., and R.D.V. wrote the article. All authors read and commented on the paper.

## Competing interests

The authors declare no competing interests.

