## Supplementary-Information for "A 6-nm ultra-photostable DNA Fluorocube for fluorescence imaging"

*for*

<u>This document includes:</u>

Supplementary Information Figures (Fig. S1 - S13)

Supplementary Information Tables (Table S1 - S10)

References for the Supplementary Information

### Supplementary Information Figures

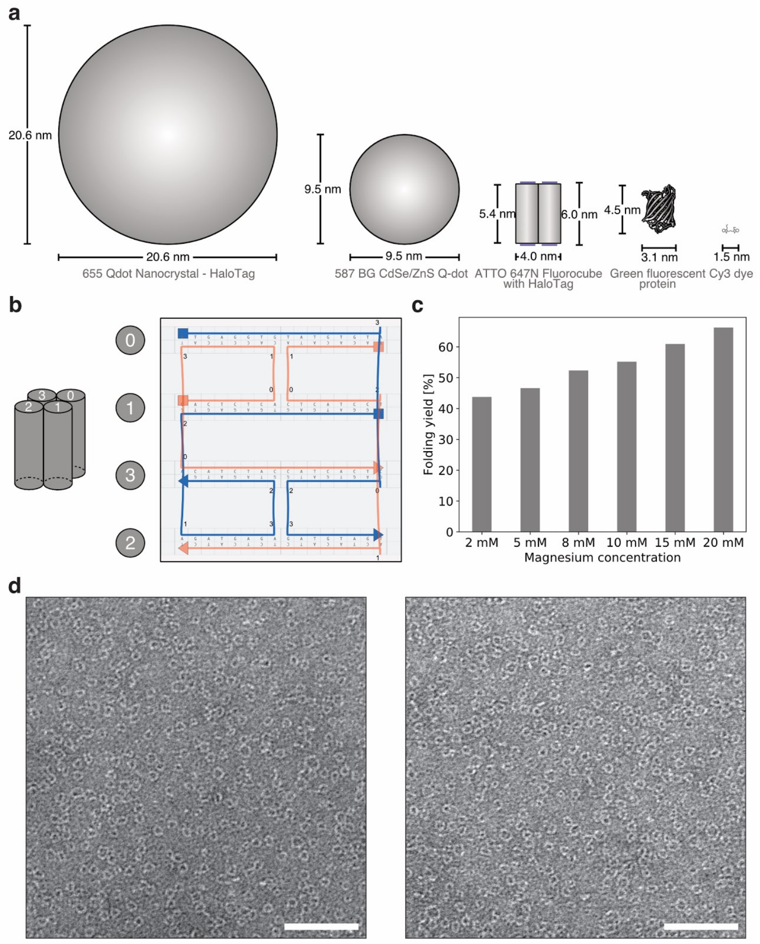


**Supplementary Figure 1 | Size, design, folding yield, and negative stain images of DNA Fluorocubes.** (**a**) Size comparison of fluorescent probes. From left to right: HALO ligand modified 655 Qdot Nanocrystal[1], benzylguanine (BG) modified 587CdSe/ZnS Q-dot[2], ATTO 647N Fluorocube with one HALO ligand[3], green fluorescent protein[4], and Cy3 dye. (**b**) Routing of four 32 bp long single-stranded DNAs (ssDNA) that are connected using crossovers. Two strands are blue and two strands are orange. The sequence is depicted below the ssDNA. A detailed list of sequences with exact modifications for all Fluorocubes used in this study is shown in **Supplementary Table 1**. The DNA routing was designed using caDNAno[5]. (**c**) Folding yield of Fluorocubes at different $MgCl_2$ concentrations from the 2% agarose gel shown in **Figure 1 c**. The folding yield was quantified using ImageJ[6]. More details about the quantification are given in the **Online Methods**. (**d**) Additional negative stain transmission electron microscopy (TEM) images of ATTO 647N Fluorocube with one biotin. Negative stain TEM images of Fluorocubes are shown in **Supplementary Figure 2**. Scale bar is 60 nm.

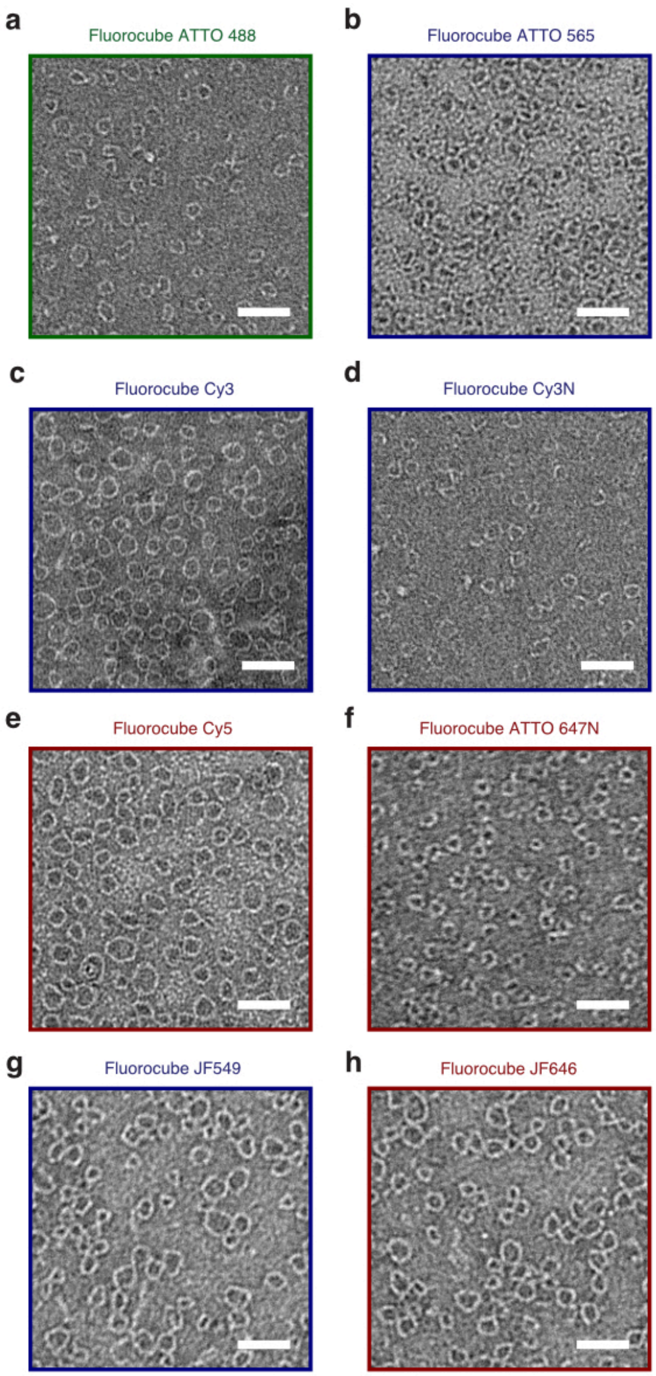


**Supplementary Figure 2 | Negative stain electron microscopy shows size variation of DNA Fluorocubes with different fluorophores.** Section of micrographs from negative stain TEM for Fluorocubes with one biotin and six fluorophores of (**a**) ATTO 488, (**b**) ATTO 565, (**c**) Cy3, (**d**) Cy3N, (**e**) Cy5, (**f**) ATTO 647N, (**g**) JF549, and (**h**) JF646. (**a-h**) Scale bar is 30 nm. "Cy3" stands for the non-sulfonated version of Cy3 whereas "Cy3N" stands for the sulfonated version of Cy3. The average diameter of Fluorocubes are given in **Supplementary Table 3 and 4**.

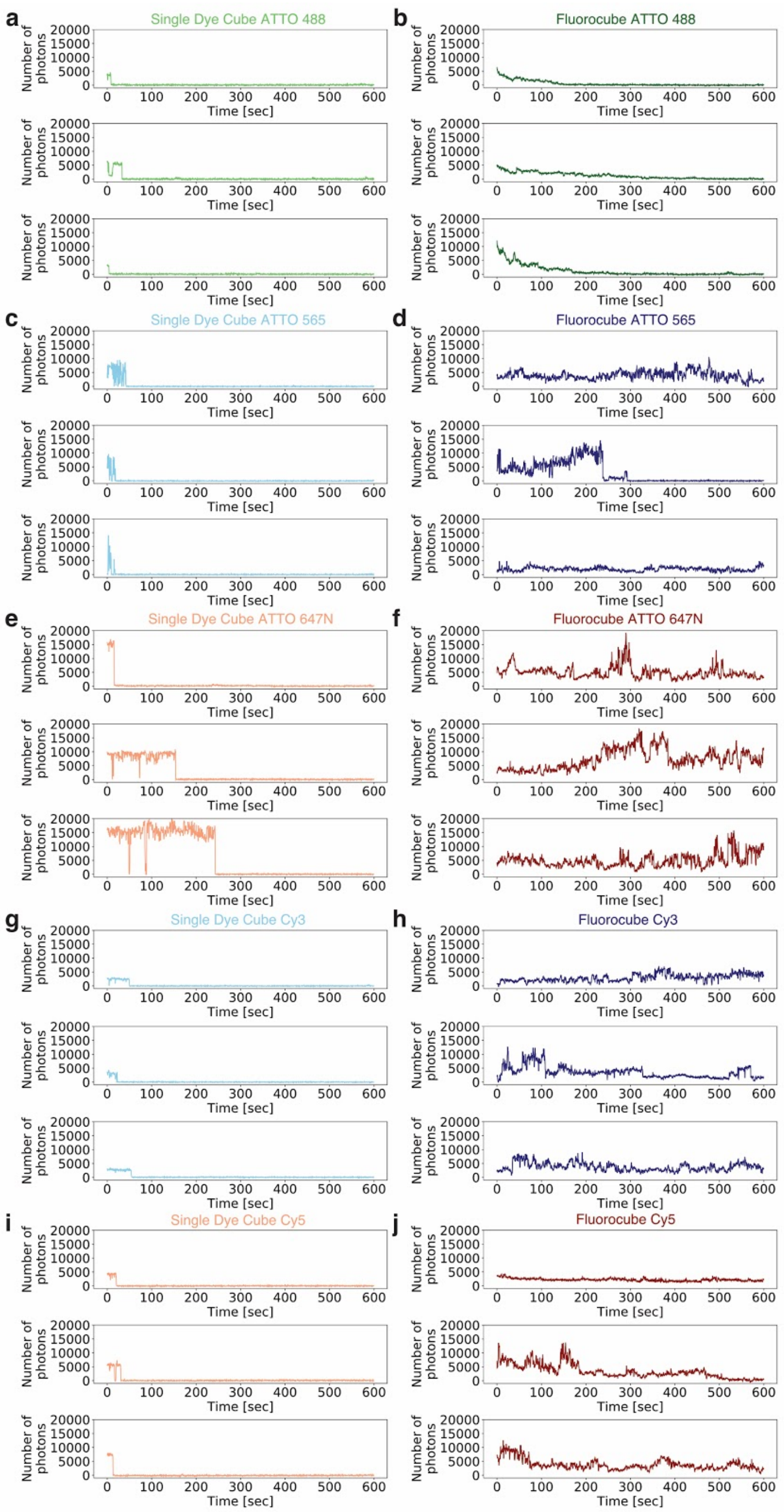
a
Single Dye Cube ATTO 488
b
Fluorocube ATTO 488
c
Single Dye Cube ATTO 565
d
Fluorocube ATTO 565
e
Single Dye Cube ATTO 647N
f
Fluorocube ATTO 647N
g
Single Dye Cube Cy3
h
Fluorocube Cy3
i
Single Dye Cube Cy5
j
Fluorocube Cy5
Number of photons
Time [sec]

**Supplementary Figure 3 | Intensity traces of DNA Fluorocubes and Single Dye Cubes with various fluorophores.** Example intensity traces of the first 600 seconds of Fluorocubes (cube with six dyes) and Single Dye Cubes (cube with one dye) used to quantify the photophysical properties shown in **Figure 1** and **Supplementary Figure 4 and 5** are presented. (**a**) Three example intensity traces for ATTO 488 Single Dye Cubes. (**b**) Three example intensity traces for ATTO 488 Fluorocubes. (**c**) Three example intensity traces for ATTO 565 Single Dye Cubes. (**d**) Three example intensity traces for ATTO 565 Fluorocubes. (**e**) Three example intensity traces for ATTO 647N Single Dye Cubes. (**f**) Three example intensity traces for ATTO 647N Fluorocubes. (**g**) Three example intensity traces for Cy3 Single Dye Cubes. (**h**) Three example intensity traces for Cy3 Fluorocubes. (**i**) Three example intensity traces for Cy5 Single Dye Cubes. (**j**) Three example intensity traces for Cy5 Fluorocubes.

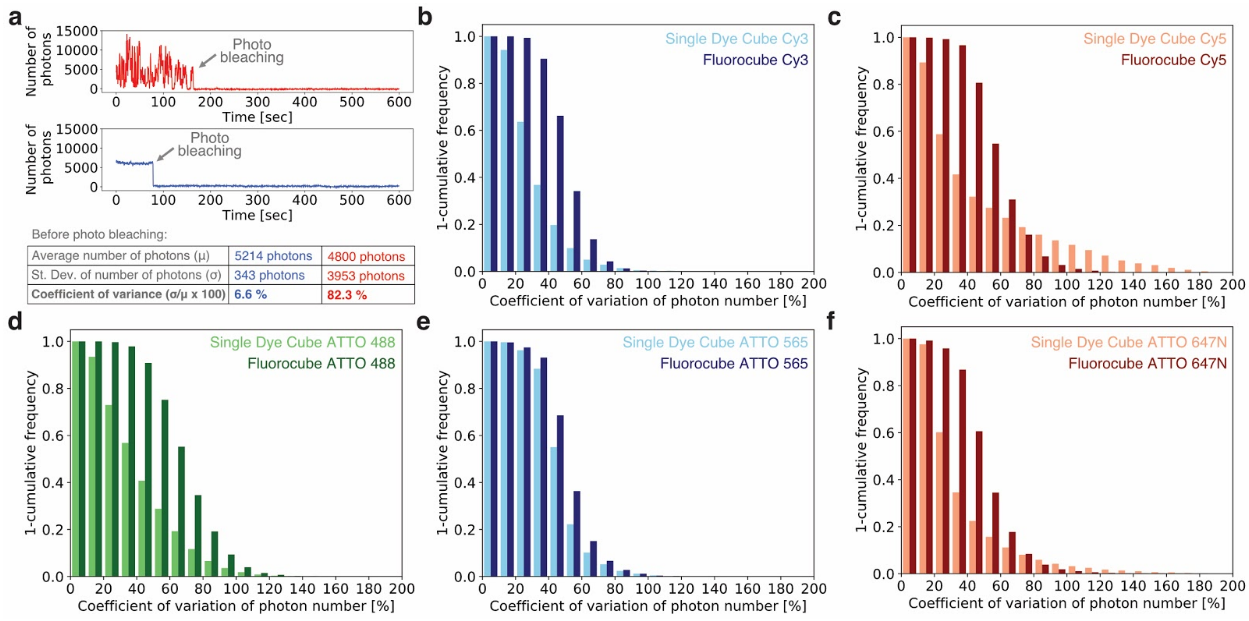


**Supplementary Figure 4 | Blinking of DNA Fluorocubes and Single Dye Cubes.** (**a**) To quantify the blinking of Fluorocubes we calculated the average number of photons and the standard deviation (St. Dev.) of number of photons per molecule up to the event of photobleaching. Using these values we determined the coefficient of variance as a measure of blinking. The top, red trace shows a Single Dye Cube that blinks a lot, whereas the bottom, blue traces shows a  Single Dye Cube with little blinking. Cumulative frequency plot for the coefficient of variance of Fluorocubes (cube with six dyes) and Single Dye Cubes (cube with one dye) with (**b**) Cy3, (**c**) Cy5, (**d**) ATTO 488, (**e**) ATTO565, and (**f**) ATTO 647N dyes. The data shown is polled from four repeats with more than 250 molecules each. Exact numbers and the standard errors of the mean of four repeats are shown in **Supplementary Table 3**.

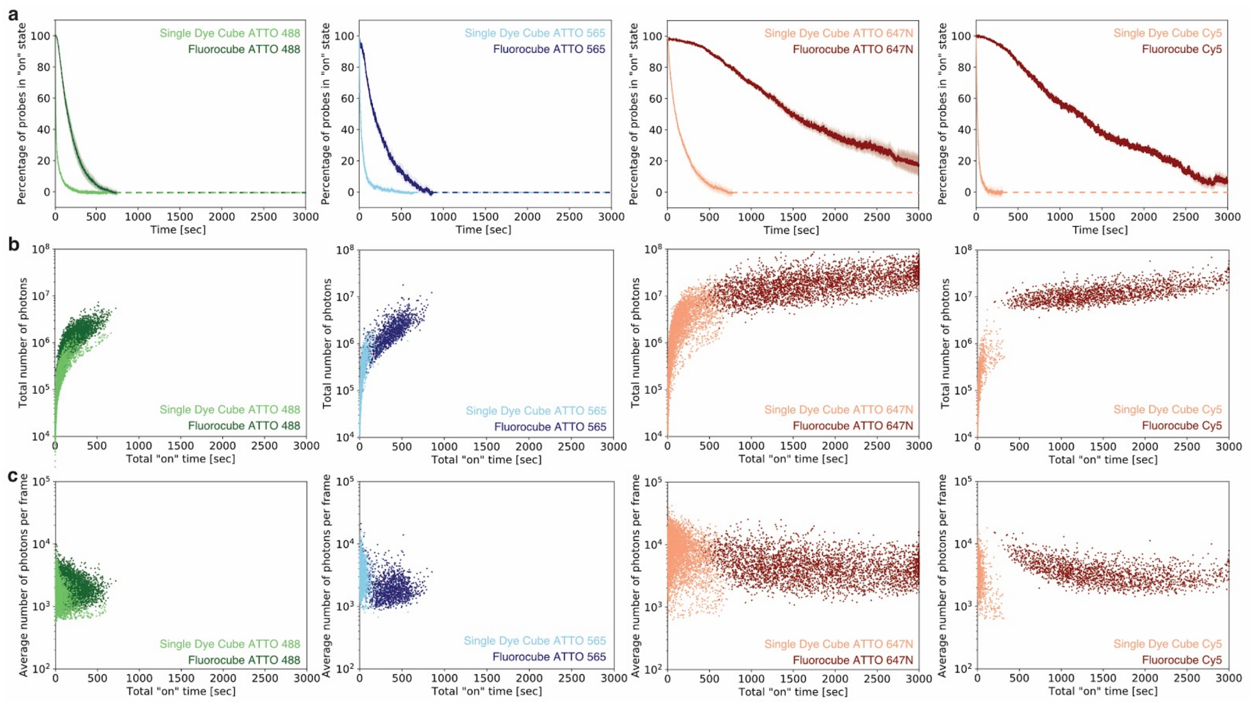


**Supplementary Figure 5 | Quantification of photophysical properties of DNA Fluorocubes and Single Dye Cubes with ATTO488, ATTO 565, ATTO 647N, and Cy5.** The experimental setup is as depicted in **Figure 1 h**. (**a**) Photostability of Fluorocubes (cube with six dyes) and Single Dye Cubes (cube with one dye) with different fluorophores. The survival rate was quantified by counting the percentage of probes in the "on" state at a given time from 0 to 3,000 seconds. Opaque color is the standard error of the mean of four repeats with more than 250 molecules each. Once all probes photobleached data analysis was terminated. This is indicated by the dashed line. (**b**) Total number of photons of Fluorocubes and Single Dye Cubes as a function of the total "on" time at the single-molecule level. (**c**) Average number of photons per frame of Fluorocubes and Single Dye Cubes as a function of the total "on" time at the single-molecule level. (**a-c**) Exact numbers are given in **Supplementary Table 3**.

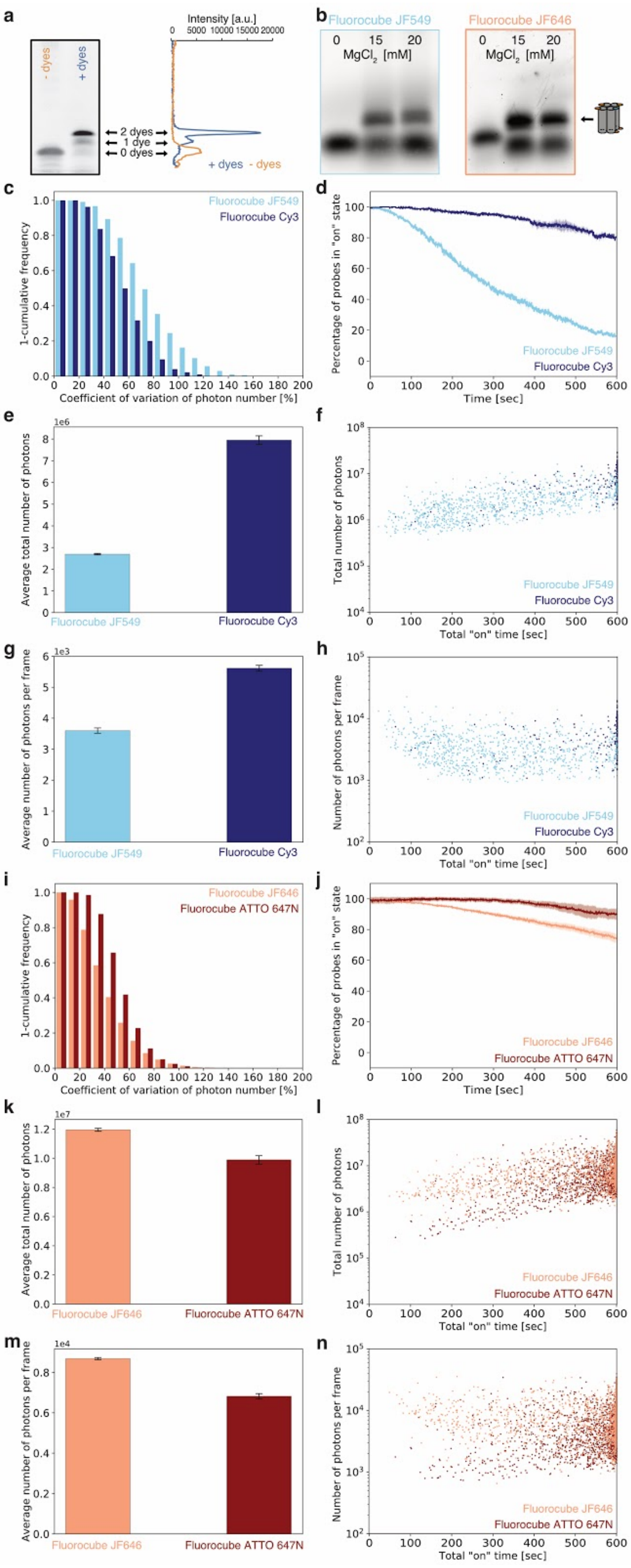

a
- dyes
+ dyes
2 dyes
1 dye
0 dyes
Intensity [a.u.]
+ dyes - dyes
b
Fluorocube JF549
Fluorocube JF646
MgCl2 [mM]
c
1-cumulative frequency
Coefficient of variation of photon number [%]
Fluorocube JF549
Fluorocube Cy3
d
Percentage of probes in "on" state
Time [sec]
e
Average total number of photons
f
Total number of photons
Total "on" time [sec]
g
Average number of photons per frame
h
Number of photons per frame
i
Fluorocube JF646
Fluorocube ATTO 647N
j
k
l
m
n

**Supplementary Figure 6 | DNA Fluorocubes with Janelia Fluorophores JF549 and JF646.** Since we noticed that the photophysical properties of Fluorocubes depend on the dye, we asked if the widely-used and very photostable Janelia Fluorophores JF549 and JF646[7] can improve the performance of Fluorocubes even further. We conjugated NHS ester Janelia Fluorophores JF549 and JF646 to amine modified oligos, assembled them into Fluorocubes and compared their photophysical properties to those of Fluorocubes with Cy3 or ATTO 647N, respectively. We again observed dye dependent behavior. While the JF646 Fluorocubes behaved equally well as the ATTO 647N Fluorocubes, the JF549 Fluorocubes performed significantly worse than the Cy3 Fluorocubes. (**a**) Labeling yield of amino group modified oligos with Janelia Fluorophores JF549 and JF646. Left PAGE column: Amino group modified oligos, which were not reacted with dyes. Right PAGE column: Amino group modified oligos, which were reacted with either Janelia Fluorophores JF549 or JF646. Labeling of ssDNA with one or two fluorophores causes a gel shift. Based on the intensity trace of a silver stained PAGE, we estimated the dual dye labeling efficiency to 76% using ImageJ[6]. More details about the labeling reaction are described in the **Online Methods**. (**b**) 2% agarose gel of Fluorocubes with Janelia Fluorophores JF549 and JF646 after thermal annealing. The four ssDNA strands are annealed at different $MgCl_2$ concentrations. Negative stain TEM images are shown in **Supplementary Figure 2 g, h**. (**c-n**) Note, that the data shown here is over 600 seconds and not over 3,000 seconds. (**c**) Cumulative frequency plot for the coefficient of variance to quantify blinking of Fluorocube with JF549 (light blue) and Fluorocube with Cy3 (dark blue). Exact numbers and the standard errors of the mean of three repeats are shown in **Supplementary Table 4**. (**d**) Photostability of Fluorocube with JF549 (light blue) and Fluorocube with Cy3 (dark blue). The survival rate was quantified by counting the percentage of probes in the "on" state at a given time from 0 to 600 seconds. Opaque color is the standard error of the mean of three repeats with more than 100 molecules each. (**e**) Histogram of the average of the total number of photons of three repeats of Fluorocube with JF549 (light blue) and Fluorocube with Cy3 (dark blue). The error bars show the standard error of the mean of three repeats with more than 100 molecules each. Exact numbers are given in **Supplementary Table 4**. Note, that the average of the total number of photons will be significantly higher than shown here because not all probes bleached within 600 seconds. (**f**) Total number of photons of Fluorocube with JF549 (light blue) and Fluorocube with Cy3 (dark blue) as a function of the total "on" time at the single-molecule level. (**g**) Histogram of the average number of photons per frame of three repeats of Fluorocube with JF549 (light blue) and Fluorocube with Cy3 (dark blue). The error bars show the standard error of the mean of three repeats with more than 100 molecules each. Exact numbers are given in **Supplementary Table 4**. (**h**) Average number of photons per frame of Fluorocube with JF549 (light blue) and Fluorocube with Cy3 (dark blue) as a function of the total “on” time at the single-molecule level. (**i-n**) Same as in **c-h** but for Fluorocube with JF646 (light red) and Fluorocube with ATTO647N (dark red).

Overall, the Fluorocube with JF549 does not perform as well as the Fluorocube with Cy3 whereas the Fluorocube with JF646 performes slightly better than the Fluorocube with ATTO647N. We note that approximately only 44% of the Fluorocubes with JF549 and JF646 have six dyes total because of the ssDNA dual labeling efficiency of 76%. Thus, with an increased labeling efficiency the Fluorocubes with JF549 and JF646 might perform even better.

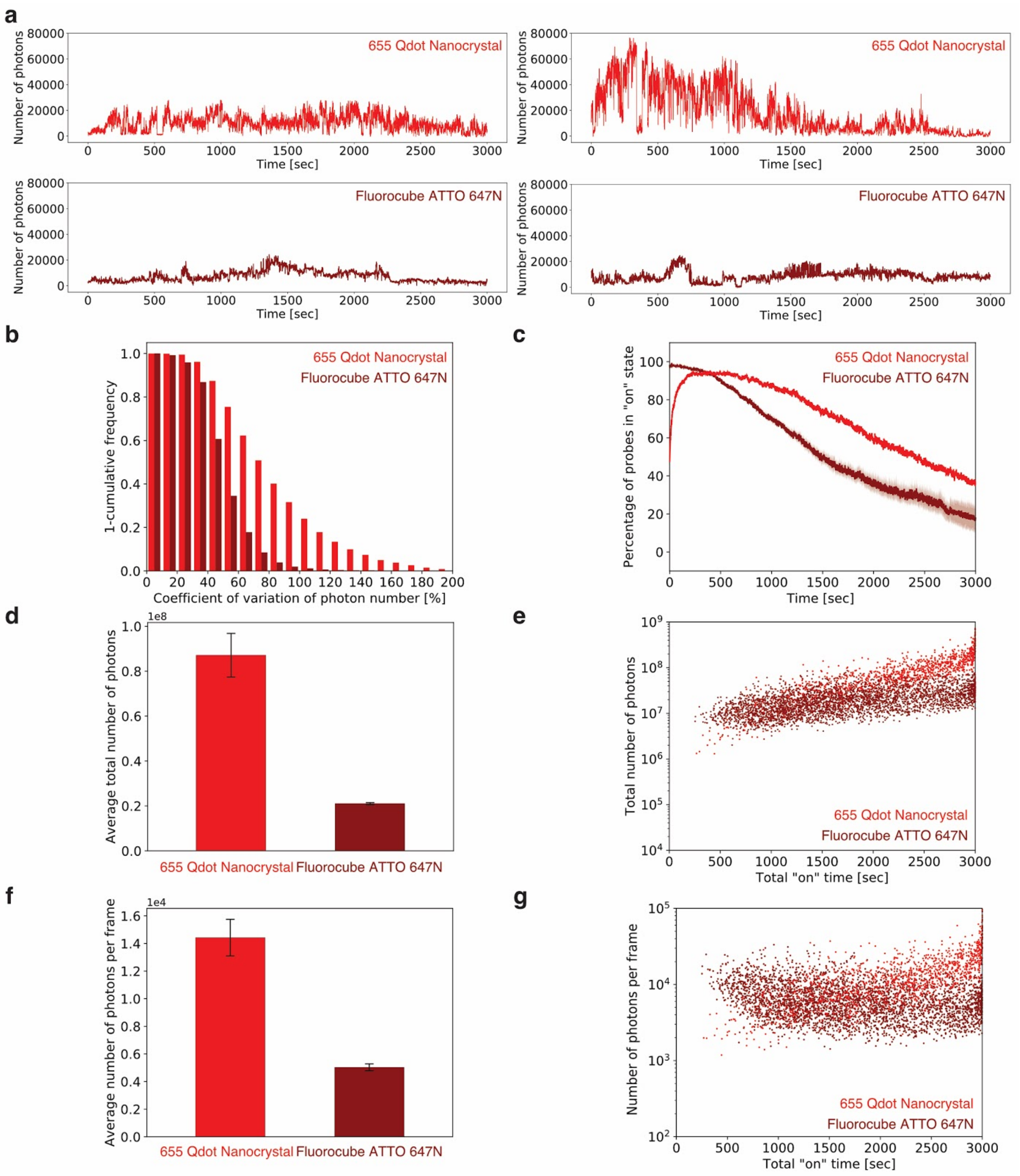


**Supplementary Figure 7 | Comparison of quantum dots with DNA Fluorocubes.** (**a**) Two example intensity traces of Streptavidin modified 655 Qdot Nanocrystal (top, light red) and ATTO 647N Fluorocube with one biotin (bottom, dark red). (**b**) Cumulative frequency plot for the coefficient of variance to quantify blinking of Streptavidin modified 655 Qdot Nanocrystal (light red) and ATTO 647N Fluorocube with one biotin (dark red). Exact numbers and the standard errors of the mean of four repeats are shown in **Supplementary Table 3**. (**c**) Photostability of Streptavidin modified 655 Qdot Nanocrystal (light red) and ATTO 647N Fluorocube with one biotin (dark red). The survival rate was quantified by counting the percentage of probes in the “on” state at any given time from 0 to 3,000 seconds. Opaque color is the standard error of the mean of four repeats with more than 500 molecules each. (**d**) Histogram of the average of the total number of photons of four repeats of Streptavidin modified 655 Qdot Nanocrystal (light red) and ATTO 647N Fluorocube with one biotin (dark red). The error bars show the standard error of the mean of four repeats with more than 500 molecules each. Exact numbers are given in **Supplementary Table 3**. Note, that the average of the total number of photons will be slightly higher than shown here because not all probes bleached within 3,000 seconds. (**e**) Total number of photons of Streptavidin modified 655 Qdot Nanocrystal (light red) and ATTO 647N Fluorocube with one biotin (dark red) as a function of the total “on” time at the single-molecule level. (**f**) Histogram of the average number of photons per frame of four experiments with Streptavidin modified 655 Qdot Nanocrystal (light red) and ATTO 647N Fluorocube with one biotin (dark red). The error bars show the standard error of the mean of four repeats with more than 500 molecules each. Exact numbers are given in **Supplementary Table 3**. (**g**) Average number of photons per frame of Streptavidin modified 655 Qdot Nanocrystal (light red) and ATTO 647N Fluorocube with one biotin (dark red) as a function of the total “on” time at the single-molecule level. For the ATTO 647N Fluorocube we are showing the same data as shown in **Supplementary Figure 4 and 5**.

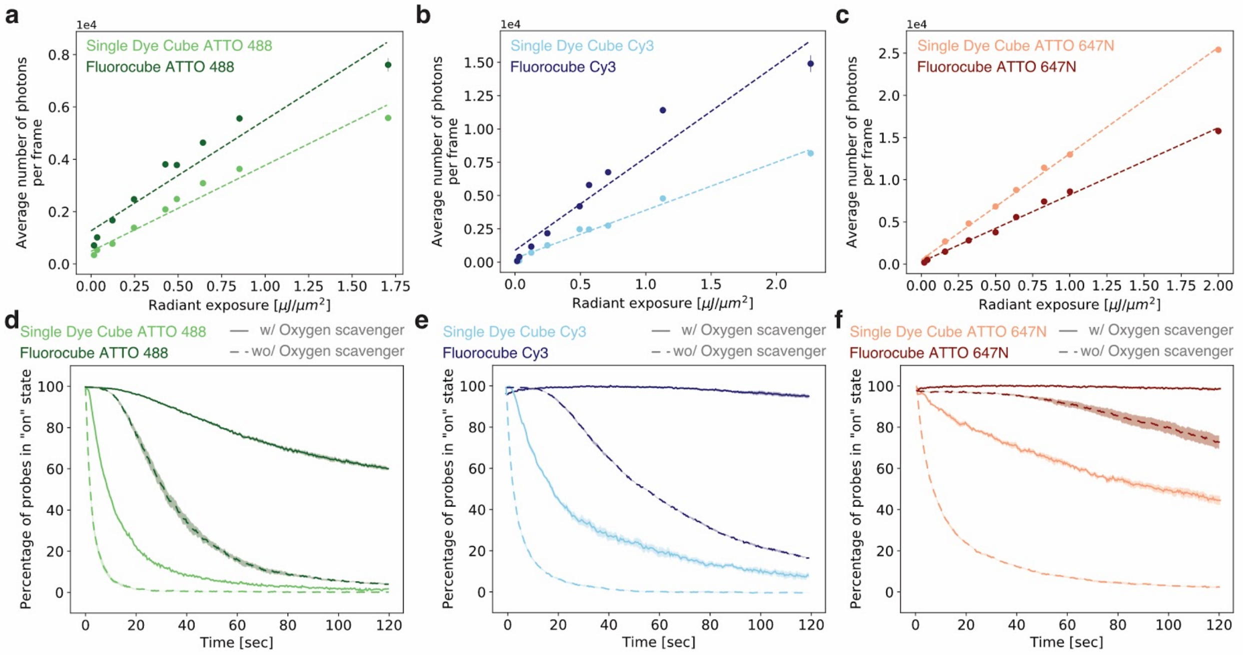


**Supplementary Figure 8 | Fluorescence intensity dependence on excitation power and effect of oxygen scavengers / triplet state quenchers.** (**a-c**) Radiant exposure was changed by reducing/increasing the laser power and/or by adjusting the exposure time for the imaging of Fluorocubes (cube with six dyes) and Single Dye Cubes (cube with one dye). A detailed list of all imaging conditions is provided in **Supplementary Table 8**. (**a**) Average number of photons per frame of three repeats as a function of radiant exposure for ATTO 488 Single Dye Cube (light green dots) and ATTO 488 Fluorocube (dark green dots). Dashed line is linear fit. Error bars show the standard error of the mean of three repeats with more than 1,000 molecules each. For most data points the error was around or less than 1% and thus the error bar is not visible. (**b**) Average number of photons per frame of three repeats as a function of radiant exposure for Cy3 Single Dye Cube (light blue dots) and Cy3 Fluorocube (dark blue dots). Dashed line is linear fit. The error bars show the standard error of the mean of three repeats with more than 200 molecules each. For most data points the error was around or less than 1% and thus the error bar is not visible. (**c**) Average number of photons per frame of three repeats as a function of radiant exposure for ATTO 647N Single Dye Cube (light red dots) and ATTO 647N Fluorocube (dark red dots). Dashed line is linear fit. The error bars show the standard error of the mean of three repeats with more than 200 molecules each. For most data points the error was around or less than 1% and thus the error bar is not visible. (**a-c**) For the probes with ATTO 488 dyes and the Cy3 Fluorocube a nonlinear dependence is observed. (**d**) Photostability of Single Dye Cubes (light green) and Fluorocubes with ATTO 488 (dark green)

with the PCA/PCD oxygen scavenging[8] and the Trolox triplet state quenching system[9] present (solid line) or absent (dashed line). (**e**) Photostability of Single Dye Cubes (light blue) and Fluorocubes with Cy3 (dark blue) with the PCA/PCD oxygen scavenging[8] and the Trolox triplet state quenching system[9] present (solid line) or absent (dashed line). (**f**) Photostability of Single Dye Cubes (light red) and Fluorocubes with ATTO 647N (dark red) with the PCA/PCD oxygen scavenging[8] and the Trolox triplet state quenching system[9] present (solid line) or absent (dashed line). (**d-f**) The survival rate was quantified by counting the percentage of probes in the "on" state at any given time from 0 to 120 seconds. Opaque color is the standard error of the mean of four repeats with more than 200 molecules each.

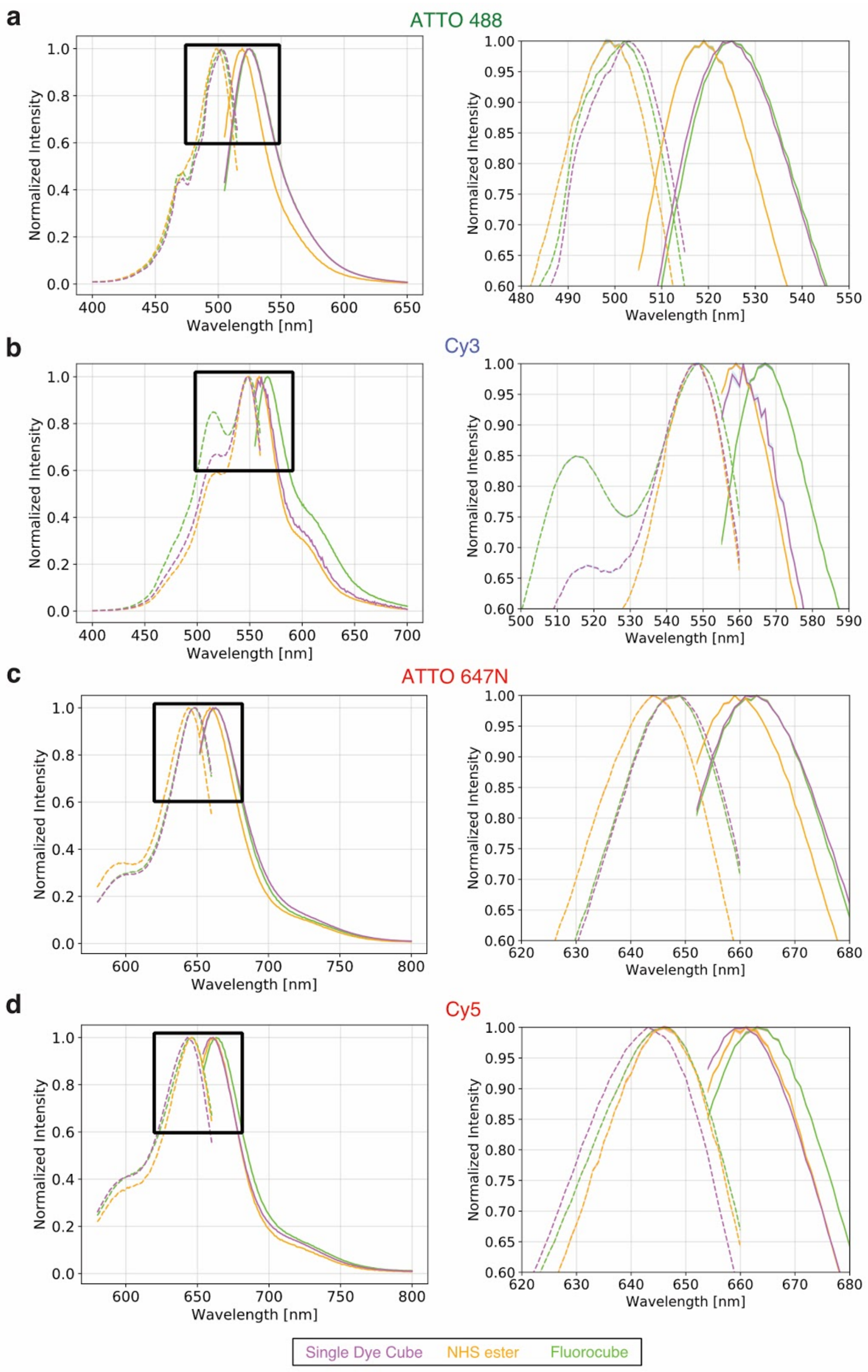


**Supplementary Figure 9 | Absorption and emission spectra of DNA Fluorocubes.** Based on the observed photophysical properties of the Fluorocubes we hypothesized that Fluorocubes may have different absorption and emission spectra than single dyes. To test this idea, we determined the absorption and emission spectra of Fluorocubes (cube with six dyes), Single

Dye Cubes (cube with one dye), and single NHS dyes. We found that the emission and absorption spectra of both ATTO dyes for the Single Dye Cube and the Fluorocube are red-shifted compared to the NHS ester dye. Thus, for the ATTO dyes, attachment to DNA seems to cause the shift rather than the close proximity of many dyes. For the cyanine dyes, we see a red-shift in the emission spectra of the Fluorocubes compared to the Single Dye Cube and NHS ester dye. Moreover, for the Cy3 Fluorocube we see a larger peak around 515 nm in the absorption spectra compared to the Single Dye Cube and NHS ester dye.

(**a-d**) Absorption (dashed line) and emission (solid line) spectra of Single Dye Cubes (purple), NHS ester dye (orange), and Fluorocubes (green) are shown for (**a**) ATTO 488, (**b**) Cy3, (**c**) ATTO 647N, and (**d**) Cy5. The area in the black box in the left panel is shown as zoom-in in the right panel. All measurements are bulk measurements. Details about instrument settings are listed in **Supplementary Tables 7**.

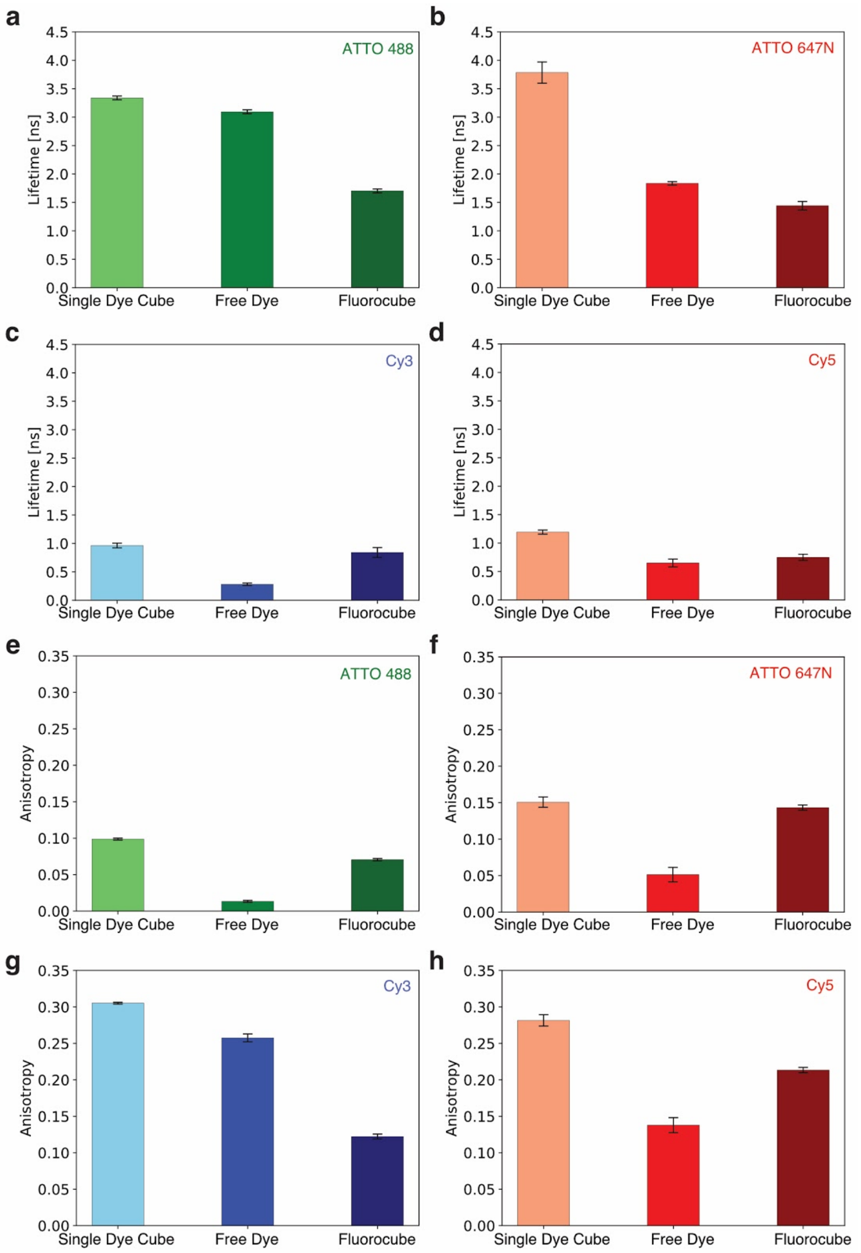


**Supplementary Figure 10 | Lifetime and anisotropy measurements of DNA Fluorocubes.** Based on the observed photophysical properties of the Fluorocubes we hypothesized that Fluorocubes may experience homo-FRET (Förster resonance energy transfer). To test this idea, we determined the lifetime and anisotropy of Fluorocubes (cube with six dyes), Single Dye Cubes (cube with one dye), and single NHS ester dyes (Free dye). We found dye specific differences between Fluorocubes and Single Dye Cubes, demonstrating that these physical parameters do not directly correlate with increased photostability of Fluorocobes.

(**a**) Fluorescence lifetime of ATTO 488 Single Dye Cubes (light green), NHS ester ATTO 488 dyes (medium green), and ATTO 488 Fluorocubes (dark green). (**b**) Fluorescence lifetime of ATTO 647N Single Dye Cubes (light red), NHS ester ATTO 647N dyes (medium red), and ATTO 647N Fluorocubes (dark red). (**c**) Fluorescence lifetime of Cy3 Single Dye Cubes (light blue), NHS ester Cy3 dyes (medium blue), and Cy3 Fluorocubes (dark blue). (**d**) Fluorescence lifetime of Cy5 Single Dye Cubes (light red), NHS ester Cy5 dyes (medium red), and Cy5 Fluorocubes (dark red). (**e**) Anisotropy measurements of ATTO 488 Single Dye Cubes (light green), NHS ester ATTO 488 dyes (medium green), and ATTO 488 Fluorocubes (dark green). (**f**) Anisotropy measurements of ATTO 647N Single Dye Cubes (light red), NHS ester ATTO 647N dyes (medium red), and ATTO 647N Fluorocubes (dark red). (**g**) Anisotropy measurements of Cy3 Single Dye Cubes (light blue), NHS ester Cy3 dyes (medium blue), and Cy3 Fluorocubes (dark blue). (**h**) Anisotropy measurements of Cy5 Single Dye Cubes (light red), NHS ester Cy5 dyes (medium red), and Cy5 Fluorocubes (dark red). (**a-h**) Error bars show standard deviation of three repeats. All measurements are bulk measurements. Details about instrument settings are listed in **Supplementary Tables 5 and 6**.

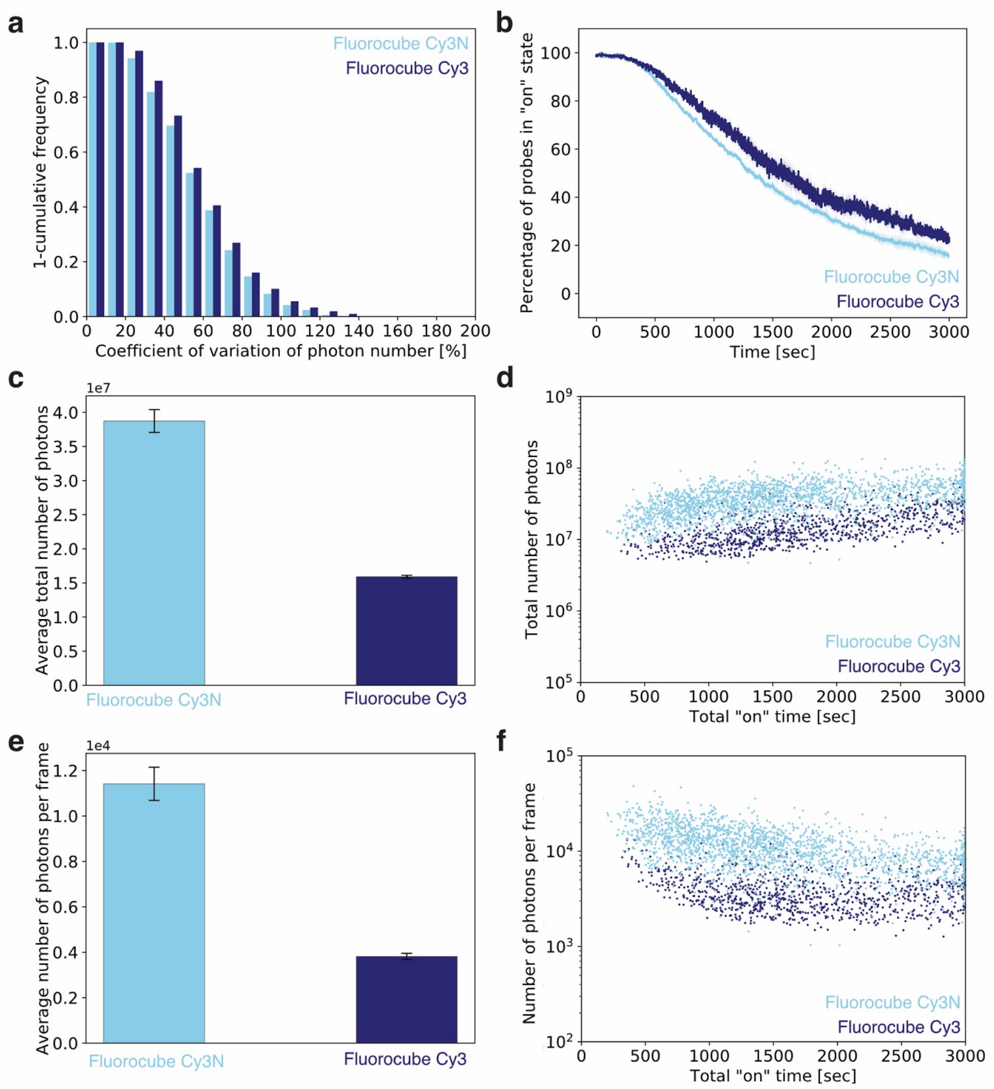


**Supplementary Figure 11 | DNA Fluorocubes with Cy3N and Cy3.** Here "Cy3" stands for the non-sulfonated version of Cy3 whereas "Cy3N" stands for the sulfonated version of Cy3. (**a**) Cumulative frequency plot for the coefficient of variance to quantify blinking of Fluorocube with Cy3N (light blue) and Fluorocube with Cy3 (dark blue). Exact numbers and the standard errors of the mean of three repeats are shown in **Supplementary Table 3**. (**b**) Photostability of Fluorocube with Cy3N (light blue) and Fluorocube with Cy3 (dark blue). The survival rate was quantified by counting the percentage of probes in the "on" state at any given time from 0 to 3000 seconds. Opaque color is the standard error of the mean of four repeats with more than 400 molecules each. (**c**) Histogram of the average of the total number of photons of four repeats of Fluorocube with Cy3N (light blue) and Fluorocube with Cy3 (dark blue). The error bars show the standard error of the mean of four repeats with more than 400 molecules each.

Exact numbers are given in **Supplementary Table 3**. Note, that the average of the total number of photons will be slightly higher than shown here because not all probes bleached within 3,000 seconds. (**d**) Total number of photons of Fluorocube with Cy3N (light blue) and Fluorocube with Cy3 (dark blue) as a function of the total "on" time at the single-molecule level. (**e**) Histogram of the average number of photons per frame of four repeats of Fluorocube with Cy3N (light blue) and Fluorocube with Cy3 (dark blue). The error bars show the standard error of the mean of four repeats with more than 400 molecules each. Exact numbers are given in **Supplementary Table 3**. (**f**) Average number of photons per frame of Fluorocube with Cy3N (light blue) and Fluorocube with Cy3 (dark blue) as a function of the total "on" time at the single-molecule level.

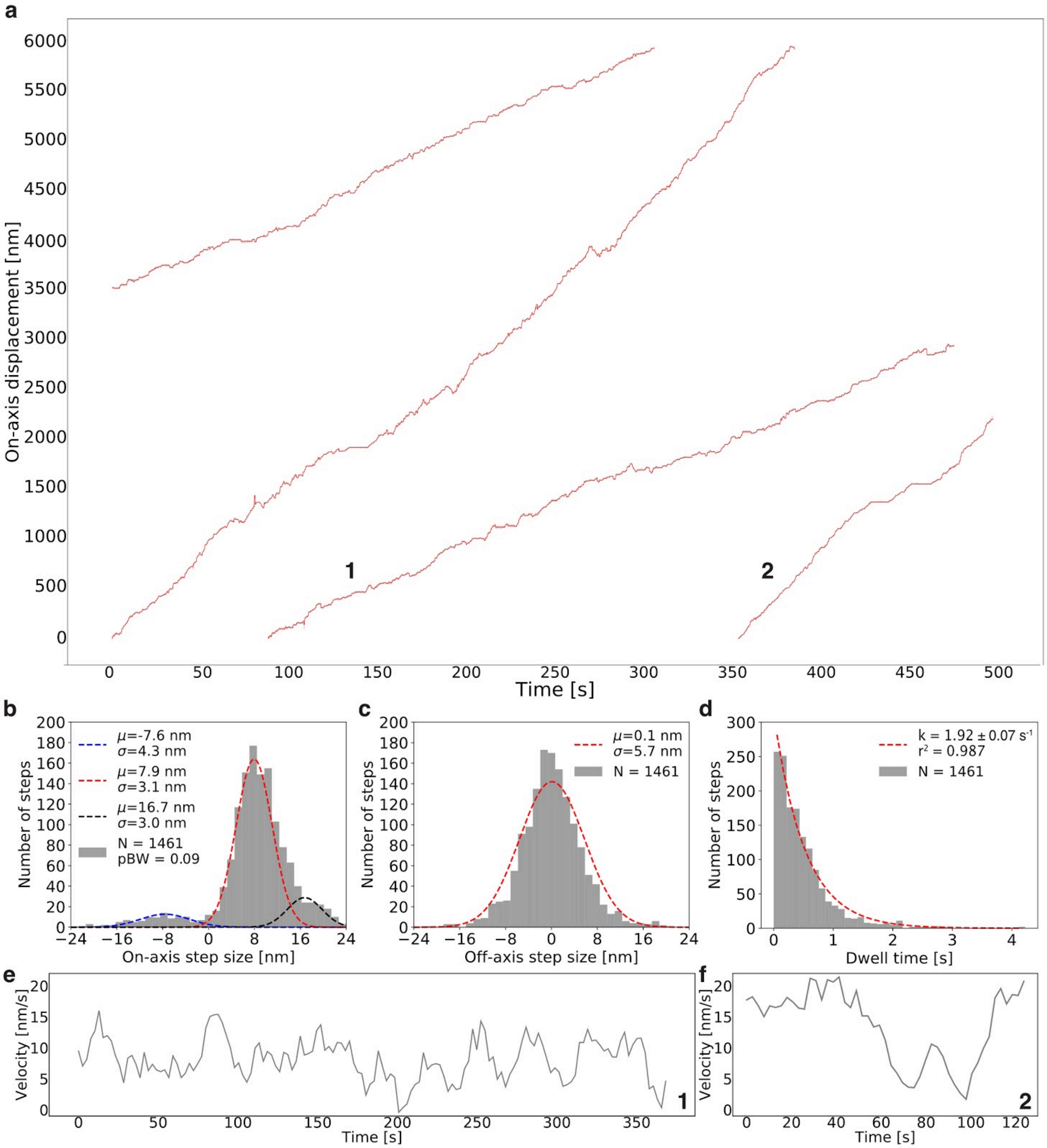


**Supplementary Figure 12 | Additional stepping data of individual kinesins.** (**a**) Raw stepping data with position versus time of four kinesins (grey dots) with detected steps (red line) along an axoneme. The opaque red line shows the standard deviation for each step. The numbers 1 and 2 indicated traces for which the velocity over time was analyzed as shown in **e** and **f**, respectively. (**b**) Histogram of the on-axis step size distribution of the combined data shown in **a**. The data was split into positive and negative steps and fit with Gaussians. For the negative steps a single Gaussian was used (blue) whereas for the positive steps two Gaussians were used (red, black). pBW is the fraction of backward steps. We mainly measured 7.9 nm forward steps which is in good agreement with previous findings for conventional kinesin 1[10] and KIF1A[11]. The 16 nm forward steps are likely two 8 nm steps which happened during a single exposure and thus could not be detected as individual steps. The observed backward steps of this artificially dimerized KIF1A[12] might be due to its properties as a monomeric motor for which backward steps have been observed[11]. In addition, the dimerization of the KIF1A motor domain using the coiled-coil of conventional kinesin 1[12], may have altered the neck linker length and alterations in neck linker length have been observed to result in few backward steps[13]. (**c**) Histogram of the off-axis step size distribution of the combined data shown in **a** fitted with a single Gaussian. The average of 0.1 nm shows that the motor moves along the axoneme with no bias to either side along the off-axis. This agrees well with previous findings that kinesin 1 prefers to move along a single protofilament without side steps[14]. (**d**) Histogram of the dwell-time distribution of the combined data shown in **a** fitted with a single exponential. The single exponential decay shows that both kinesin heads step at a similar rate, which in this case is limited by the ATP concentration[15]. (**e, f**) Velocity over time for stepping traces of a single kinesin as shown in **a**. The grey line shows a moving average of velocity binned into 15.6 sec (for details see **Online Methods**). The numbers 1 and 2 indicate which traces in **a** where chosen.

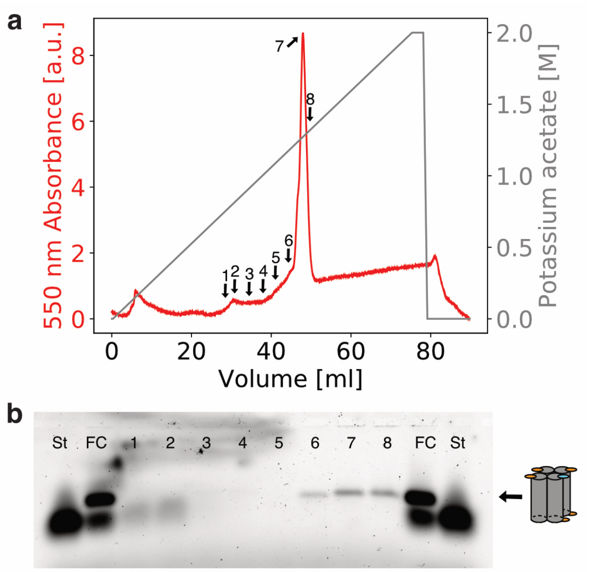


**Supplementary Figure 13 | Ion exchange chromatography separates DNA Fluorocubes from excess oligos.** (**a**) 550 nm absorbance trace (red) from ion exchange chromatography with Cy3 Fluorocubes with one biotin. The grey trace shows the linear increase of potassium acetate from 0 M to 2 M over a volume of 80 ml. The arrows with numbers indicate fractions that were run on a 2% agarose gel to evaluate elution fractions. (**b**) 2% agarose gel with elution fractions from ion exchange chromatography (1-8) as shown in **a**. “St” refers to a single of the four ssDNA (FC_St_02) used to fold the Fluorocubes (**Supplementary Table 1**). “FC” refers to a Cy3 Fluorocube with one biotin that was not purified by ion exchange chromatography. Based on the agarose gel we can conclude that most of the Fluorocubes elute at approximately 1.1 M potassium acetate. Moreover, oligos that were not incorporated into a structure (Fluorocube) are clearly separated from fully folded Fluorocubes. We verified Fluorocube integrity by negative stain electron microscopy after ion exchange chromatography. Taken together, ion exchange chromatography can be used to further purify Fluorocubes.

### Supplementary Information Tables

| Name | Sequence and modification | Vendor |
|---|---|---|
| FC_SC_01_NH2 | /5AmMC6/ATGAGGTGTATGTGTAGAGTGATGGATGTAGT/3AmMO/ | IDT |
| FC_SC_02_NH2 | /5AmMC6/AGGATGAGTGAGAGTGAGATGAGAGTAGATGT/3AmMO/ | IDT |
| FC_St_02_NH2 | /5AmMC6/CACTCTCACACCTCATACATCTACCATCACTC/3AmMO/ | IDT |
| FC_SC_01_ATTO488 | /5ATTO488N/ATGAGGTGTATGTGTAGAGTGATGGATGTAGT/3ATTO488N/ | IDT |
| FC_SC_02_ATTO488 | /5ATTO488N/AGGATGAGTGAGAGTGAGATGAGAGTAGATGT/3ATTO488N/ | IDT |
| FC_St_02_ATTO488 | /5ATTO488N/CACTCTCACACCTCATACATCTACCATCACTC/3ATTO488N/ | IDT |
| FC_SC_01_ATTO565 | /5ATTO565N/ATGAGGTGTATGTGTAGAGTGATGGATGTAGT/3ATTO565N/ | IDT |
| FC_SC_02_ATTO565 | /5ATTO565N/AGGATGAGTGAGAGTGAGATGAGAGTAGATGT/3ATTO565N/ | IDT |
| FC_St_02_ATTO565 | /5ATTO565N/CACTCTCACACCTCATACATCTACCATCACTC/3ATTO565N/ | IDT |
| FC_SC_01_ATTO647N | /5ATTO647NN/ATGAGGTGTATGTGTAGAGTGATGGATGTAGT/3ATTO647NN/ | IDT |
| FC_SC_02_ATTO647N | /5ATTO647NN/AGGATGAGTGAGAGTGAGATGAGAGTAGATGT/3ATTO647NN/ | IDT |
| FC_St_02_ATTO647N | /5ATTO647NN/CACTCTCACACCTCATACATCTACCATCACTC/3ATTO647NN/ | IDT |
| FC_SC_01_Cy3 | /5Cy3/ATGAGGTGTATGTGTAGAGTGATGGATGTAGT/3Cy3Sp/ | IDT |
| FC_SC_02_Cy3 | /5Cy3/AGGATGAGTGAGAGTGAGATGAGAGTAGATGT/3Cy3Sp/ | IDT |
| FC_St_02_Cy3 | /5Cy3/CACTCTCACACCTCATACATCTACCATCACTC/3Cy3Sp/ | IDT |
| FC_SC_01_Cy3N | /5Cy3N/ATGAGGTGTATGTGTAGAGTGATGGATGTAGT/3Cy3N/ | IDT |
| FC_SC_02_Cy3N | /5Cy3N/AGGATGAGTGAGAGTGAGATGAGAGTAGATGT/3Cy3N/ | IDT |
| FC_St_02_Cy3N | /5Cy3N/CACTCTCACACCTCATACATCTACCATCACTC/3Cy3N/ | IDT |
| FC_SC_01_Cy5 | /5Cy5/ATGAGGTGTATGTGTAGAGTGATGGATGTAGT/3Cy5Sp/ | IDT |
| FC_SC_02_Cy5 | /5Cy5/AGGATGAGTGAGAGTGAGATGAGAGTAGATGT/3Cy5Sp/ | IDT |
| FC_St_02_Cy5 | /5Cy5/CACTCTCACACCTCATACATCTACCATCACTC/3Cy5Sp/ | IDT |
| FC_St_01_Biotin | /5Biosg/TACACATACTCATCCTACTACATCTCTCATCT | IDT |
| FC_St_01_HALO | Halotag Ligand (O2) TACACATACTCATCCTACTACATCTCTCATCT | Biomers |
| SD_SC_01_UN | ATGAGGTGTATGTGTAGAGTGATGGATGTAGT | IDT |
| SD_SC_02_UN | AGGATGAGTGAGAGTGAGATGAGAGTAGATGT | IDT |
| SD_St_02_ATTO488 | /5ATTO488N/CACTCTCACACCTCATACATCTACCATCACTC | IDT |
| SD_St_02_ATTO565 | /5ATTO565N/CACTCTCACACCTCATACATCTACCATCACTC | IDT |
| SD_St_02_ATTO647N | /5ATTO647NN/CACTCTCACACCTCATACATCTACCATCACTC | IDT |
| SD_St_02_Cy3 | /5Cy3/CACTCTCACACCTCATACATCTACCATCACTC | IDT |
| SD_St_02_Cy5 | /5Cy5/CACTCTCACACCTCATACATCTACCATCACTC | IDT |

**Supplementary Table 1 | Sequences and corresponding modifications of all oligonucleotides used in this study.** Nomenclature: "FC" stands for a single-stranded DNA (ssDNA) strand that was used to fold Fluorocubes (cube with six dyes). "SD" stands for a ssDNA strand that was used to fold Single Dye Cubes (cube with one dye). "NH2" stands for amino group modification. "UN" stands for unlabeled oligonucleotide.

For each Fluorocube and Single Dye Cube four ssDNA strands are required: SC_01, SC_02, St_01, and St_02. Both Single Dye Cubes (light gray part of the table) and Fluorocubes (white part of the table) need a functionalized oligo strand, which is always St_01 (dark gray part of the table), and either has a biotin or a HALO-ligand[3]. Single Dye Cubes, irrespective of the dye, always require two unlabeled strands "SD_SC_01_UN" and "SD_SC_02_UN" as well as one strand with a single dye "SD_St_02". Thus, for example, to assemble a Single Dye Cube with ATTO488 and biotin, one uses: SD_SC_01_UN, SD_SC_02_UN, FC_St_01_Biotin, and SD_St_02_ATTO488. For a Fluorocube with biotin one uses: FC_SC_01_ATTO647N, FC_SC_02_ATTO647N, FC_St_01_Biotin, and FC_St_02_ATTO647N. Here "Cy3" stands for the non-sulfonated version of Cy3 whereas "Cy3N" stands for the sulfonated version of Cy3. A detailed overview of strand combinations used for Single Dye Cubes and Fluorocubes can be found in **Supplementary Table 2**. Note that amino group modified oligos were used to label with either JF549 or JF646.

| Fluorocube / Single Dye Cube name | Oligonucleotide stands used |
|---|---|
| FC_ATTO488_Biotin | FC_SC_01_ATTO488, FC_SC_02_ATTO488, FC_St_01_Biotin, FC_St_02_ATTO488 |
| FC_ATTO565_Biotin | FC_SC_01_ATTO565, FC_SC_02_ATTO565, FC_St_01_Biotin, FC_St_02_ATTO565 |
| FC_ATTO647N_Biotin | FC_SC_01_ATTO647N, FC_SC_02_ATTO647N, FC_St_01_Biotin, FC_St_02_ATTO647N |
| FC_ATTO647N_HALO | FC_SC_01_ATTO647N, FC_SC_02_ATTO647N, FC_St_01_HALO, FC_St_02_ATTO647N |
| FC_Cy3_Biotin | FC_SC_01_Cy3, FC_SC_02_Cy3, FC_St_01_Biotin, FC_St_02_Cy3 |
| FC_Cy3N_Biotin | FC_SC_01_Cy3N, FC_SC_02_Cy3N, FC_St_01_Biotin, FC_St_02_Cy3N |
| FC_Cy5_Biotin | FC_SC_01_Cy5, FC_SC_02_Cy5, FC_St_01_Biotin, FC_St_02_Cy5 |
| FC_JF549_Biotin | FC_SC_01_NH2, FC_SC_02_NH2, FC_St_01_Biotin, FC_St_02_NH2 |
| FC_JF646_Biotin | FC_SC_01_NH2, FC_SC_02_NH2, FC_St_01_Biotin, FC_St_02_NH2 |
| SD_ATTO488_Biotin | SD_SC_01_UN, SD_SC_02_UN, FC_St_01_Biotin, SD_St_02_ATTO488 |
| SD_ATTO565_Biotin | SD_SC_01_UN, SD_SC_02_UN, FC_St_01_Biotin, SD_St_02_ATTO565 |
| SD_ATTO647N_Biotin | SD_SC_01_UN, SD_SC_02_UN, FC_St_01_Biotin, SD_St_02_ATTO647N |
| SD_Cy3_Biotin | SD_SC_01_UN, SD_SC_02_UN, FC_St_01_Biotin, SD_St_02_Cy3 |
| SD_Cy5_Biotin | SD_SC_01_UN, SD_SC_02_UN, FC_St_01_Biotin, SD_St_02_Cy5 |

**Supplementary Table 2 | Combination of oligonucleotide strands used to assemble all DNA Fluorocubes and Single Dye Cubes used in this study.** Nomenclature: “FC” stands for Fluorocubes with six dyes and “SD” stands for Single Dye Cubes. The exact sequences with modifications are given in **Supplementary Table 1**. Here “Cy3” stands for the non-sulfonated version of Cy3 whereas “Cy3N” stands for the sulfonated version of Cy3. Note, that for Fluorocubes with either Janelia Fluorophores JF549 or JF646 amino group modified oligos have been used which were labeled with either dye as described in the **Online Methods** and shown in **Supplementary Figure 6**. A protocol for annealing the oligonucleotide strands to fold DNA Fluorocubes and Single Dye Cubes is given in the **Online Methods**.

| | Photo-bleaching half-life time in seconds | Total number of photons ($10^6$) | Average number of photons per frame ($10^3$) | Blinking - Coefficient of variation of photon number | Average outside diameter of Fluorocube by negative stain in nanometer | Charge per dye |
|---|---|---|---|---|---|---|
| **Single Dye Cube (ATTO 488) n = 4390** | 8 ± 1 | 0.19 ± 0.02 | 1.93 ± 0.04 | 39.8 ± 1.1 | n.m. | -1 |
| **Fluorocube (ATTO 488) n = 7916** | 149 ± 3 | 0.83 ± 0.07 | 2.01 ± 0.06 | 63.9 ± 0.2 | 6.75 ± 1.31 (n = 141) | -1 |
| **Single Dye Cube (ATTO 565) n = 2349** | 15 ± 2 | 0.35 ± 0.05 | 3.55 ± 0.09 | 43.0 ± 0.2 | n.m. | 0 |
| **Fluorocube (ATTO 565) n = 1368** | 181 ± 4 | 2.49 ± 0.08 | 2.51 ± 0.10 | 46.4 ± 1.1 | 7.79 ± 1.28 (n = 98) | 0 |
| **Single Dye Cube (ATTO 647N) n = 4359** | 98 ± 3 | 2.94 ± 0.02 | 8.49 ± 0.35 | 33.3 ± 1.8 | n.m. | +1 |
| **Fluorocube (ATTO 647N) n = 3495** | 1532 ± 9 | 21.04 ± 0.44 | 5.03 ± 0.25 | 46.1 ± 0.4 | 6.24 ± 1.17 (n = 189) | +1 |
| **Single Dye Cube (Cy3) n = 2260** | 24 ± 2 | 0.39 ± 0.01 | 2.79 ± 0.04 | 28.3 ± 0.2 | n.m. | +1 |
| **Fluorocube (Cy3) n = 2290** | 1389 ± 6 | 15.71 ± 0.37 | 3.96 ± 0.10 | 45.8 ± 0.7 | 9.18 ± 2.24 (n = 146) | +1 |
| **Fluorocube (Cy3N) n = 1966** | 1283 ± 7 | 38.37 ± 0.57 | 11.14 ± 0.42 | 44.1 ± 0.6 | 6.80 ± 1.41 (n = 128) | -1 |
| **Single Dye Cube (Cy5) n = 1207** | 17 ± 1 | 0.41 ± 0.08 | 3.95 ± 0.18 | 42.6 ± 2.1 | n.m. | +1 |
| **Fluorocube (Cy5) n = 1901** | 1244 ± 5 | 12.82 ± 0.62 | 3.72 ± 0.14 | 54.1 ± 0.1 | 9.96 ± 2.33 (n = 119) | +1 |
| **655 Qdot n = 1341** | 2385 ± 11 | 87.12 ± 0.97 | 14.43 ± 1.32 | 100.2 ± 5.2 | n.m. | N.A. |

**Supplementary Table 3 | Photophysical properties of quantum dots, DNA Fluorocubes, and Single Dye Cubes with different fluorophores.** Here “Single Dye Cube” refers to a cube with a single dye whereas “Fluorocube” refers to a cube with six dyes. In all cases the probes had a single biotin. Here “Cy3” stands for the non-sulfonated version of Cy3 whereas “Cy3N” stands for the sulfonated version of Cy3. The values listed for “Cy3” are from the data shown in **Figure 1** and not from **Supplementary Figure 11**, which were very similar. Note, that the average of the total number of photons shown here is an underestimate because not all probes bleached within 3,000 seconds. Errors (±) are the standard error of the mean of four repeats. “n.m.” is not measured and “N.A.” is not available.

| | Total number of photons ($10^6$) | Average number of photons per frame ($10^3$) | Blinking - Coefficient of variation of photon number | Average outside diameter of Fluorocube by negative stain in nanometer | Charge per dye |
|---|---|---|---|---|---|
| **Fluorocube (Cy3) n = 325** | 7.95 ± 0.19 | 5.61 ± 0.09 | 51.9 ± 0.1 | See Supplementary Table 3 | +1 |
| **Fluorocube (JF549) n = 918** | 2.69 ± 0.03 | 3.60 ± 0.09 | 72.1 ± 1.9 | 7.61 ± 1.72 (n = 133) | 0 |
| **Fluorocube (ATTO 647N) n = 3025** | 9.89 ± 0.29 | 6.82 ± 0.13 | 46.6 ± 2.3 | See Supplementary Table 3 | +1 |
| **Fluorocube (JF646) n = 2518** | 11.96 ± 0.10 | 8.68 ± 0.05 | 37.6 ± 1.4 | 7.24 ± 1.70 (n = 103) | 0 |

**Supplementary Table 4 | Photophysical properties of DNA Fluorocubes with Janelia fluorophores.** Here “Fluorocube” refers to a cube with six dyes. In all cases the probes had a single biotin. Note, that the average total number of photons shown here is an underestimate because most probes did not bleach within 600 seconds. Errors (±) are the standard error of the mean of three repeats.

| | Iterations | Excitation Wavelength (fixed) | Emission Wavelength (fixed) |
|---|---|---|---|
| **ATTO 488** | 24 | 500 nm | 520 nm |
| **ATTO 647N** | 24 | 647 nm | 665 nm |
| **Cy3** | 24 | 550 nm | 565 nm |
| **Cy5** | 24 | 649 nm | 665 nm |

**Supplementary Table 5 | Settings for anisotropy measurements.** These settings were used for all samples, Fluorocubes (cube with six dyes), Single Dye Cubes (cube with one dye), and single NHS ester dyes. All measurements are bulk measurements.

| | Excitation Wavelength (fixed) | Emission Longpass filter | Modulation frequencies |
|---|---|---|---|
| **ATTO 488** | 470 nm | 500 nm | 400,000 Hz, 797,895 Hz, 1,591,590 Hz, 3,174,802 Hz, 6,332,894 Hz, 12,632,455 Hz, 25,198,421 Hz, 50,264,213 Hz, 100,263,864 Hz, and 200,000,000 Hz |
| **ATTO 647N** | 620 nm | 650 nm | See ATTO 488 |
| **Cy3** | 475 nm | 520 nm | See ATTO 488 |
| **Cy5** | 620 nm | 650 nm | See ATTO 488 |

**Supplementary Table 6 | Settings for fluorescence lifetime measurements.** These settings were used for all samples, Fluorocubes (cube with six dyes), Single Dye Cubes (cube with one dye), and single NHS ester dyes. All measurements are bulk measurements.

| | Absorption spectra - Excitation Wavelength | Absorption spectra - Emission Wavelength (fixed) | Emission spectra - Excitation Wavelength (fixed) | Emission spectra - Emission Wavelength |
|---|---|---|---|---|
| **ATTO 488** | 400 to 515 nm | 520 nm | 500 nm | 505 to 650 nm |
| **ATTO 647N** | 580 to 660 nm | 665 nm | 647 nm | 652 to 800 nm |
| **Cy3** | 400 to 560 nm | 565 nm | 550 nm | 555 to 700 nm |
| **Cy5** | 580 to 660 nm | 665 nm | 649 nm | 654 to 800 nm |

**Supplementary Table 7 | Settings for absorption and emission spectra measurements.** These settings were used for all samples, Fluorocubes (cube with six dyes), Single Dye Cubes (cube with one dye), and single NHS ester dyes. We used an interval of 1 nm between measurements. All measurements are bulk measurements.

| Laser power 488 nm [mW] | Laser power 561 nm [mW] | Laser power 640 nm [mW] | Exposure time [ms] | Radiant exposure 488 nm [µJ/µm$^2$] | Radiant exposure 561 nm [µJ/µm$^2$] | Radiant exposure 640 nm [µJ/µm$^2$] |
|---|---|---|---|---|---|---|
| 0.31 | 0.29 | 0.36 | 100 | 0.01 | 0.01 | 0.01 |
| 0.31 | 0.29 | 0.36 | 200 | 0.02 | 0.02 | 0.02 |
| 0.31 | 0.29 | 0.36 | 400 | 0.03 | 0.03 | 0.04 |
| 4.44 | 4.45 | 5.75 | 100 | 0.12 | 0.12 | 0.16 |
| 4.44 | 4.45 | 5.75 | 200 | 0.25 | 0.25 | 0.32 |
| 11.58 | 12.82 | 14.89 | 100 | 0.32 | 0.36 | 0.41 |
| 11.58 | 12.82 | 14.89 | 200 | 0.64 | 0.71 | 0.83 |
| 11.58 | 12.82 | 14.89 | 400 | 1.29 | 1.42 | 1.65 |
| 15.36 | 20.36 | 18.01 | 100 | 0.43 | 0.57 | 0.50 |
| 15.36 | 20.36 | 18.01 | 200 | 0.85 | 1.13 | 1.00 |
| 15.36 | 20.36 | 18.01 | 400 | 1.71 | 2.26 | 2.00 |

**Supplementary Table 8 | Calculation of radiant exposures as used in Supplementary Figure 8.** Laser power was measured after the objective. The field of illumination has a 2D Gaussian shape (reflecting the Gaussian shape of the laser beam), causing the radiant exposure to vary over the field of view. Here, we calculated an average radiant exposure by assuming a field of illumination of 60 µm by 60 µm.

| Figure | Dataset | Number of frames to check |
|---|---|---|
| Fig. 1, Supp. Fig. 4, Supp. Fig. 5 | Single Dye Cube (ATTO 488) | 10 |
| Fig. 1, Supp. Fig. 4, Supp. Fig. 5 | Fluorocube (ATTO 488) | 1,000 |
| Fig. 1, Supp. Fig. 4 | Single Dye Cube (Cy3) | 10 |
| Fig. 1, Supp. Fig. 4, Supp. Fig. 6, Supp. Fig. 11 | Fluorocube (Cy3) | 7,500 |
| Fig. 1, Supp. Fig. 4, Supp. Fig. 5 | Single Dye Cube (ATTO 647N) | 10 |
| Fig. 1, Supp. Fig. 4, Supp. Fig. 5, Supp. Fig. 6, Supp. Fig. 7 | Fluorocube (ATTO 647N) | 7,500 |
| Fig. 1, Supp. Fig. 4, Supp. Fig. 5 | Single Dye Cube (ATTO 565) | 10 |
| Fig. 1, Supp. Fig. 4, Supp. Fig. 5 | Fluorocube (ATTO 565) | 1,000 |
| Fig. 1, Supp. Fig. 4, Supp. Fig. 5 | Single Dye Cube (Cy5) | 10 |
| Fig. 1, Supp. Fig. 4, Supp. Fig. 5 | Fluorocube (Cy5) | 7,500 |
| Fig. 1, Supp. Fig. 6 | Fluorocube (JF 549) | 1,000 |
| Fig. 1, Supp. Fig. 6 | Fluorocube (JF 646) | 7,500 |
| Supp. Fig. 7 | Q-dot | 7,500 |
| Fig. 1, Supp. Fig. 11 | Fluorocube (Cy3N) | 7,500 |

**Supplementary Table 9 | Number of first frames to check for Spot Intensity Analysis Plug in.** The “Spot Intensity Plugin” finds spot coordinates by averaging a user-definable number of frames and detecting local maxima within this averaged image. Intensities at these coordinates are then measured in the complete data set. The number of frames to check varies for each sample depending on how fast they bleach. If for example a molecule bleaches early (as for the Single Dye Cubes), many dark frames will be used for the average and very few molecules will be found since the average intensity will be lower than the threshold (Noise). In a similar way, if too few frames are used and a molecule gets brighter over time (as for the Fluorocubes), then it may not be considered. Thus, we typically used only a few frames for the Single Dye Cube samples and more frames for the Fluorocube samples. However, this also depended on how fast the Fluorocubes bleached. “Single Dye Cube” refers to a cube with a single dye whereas

“Fluorocube” refers to a cube with six dyes. “Cy3” stands for the non-sulfonated version of Cy3 whereas “Cy3N” stands for the sulfonated version of Cy3.

| **Imaging Parameters** | |
|---|---|
| Photon conversion factor | 1.84 |
| Linear (EM) gain | 1.0 |
| Pixel size [nm] | 159.0 |
| Time interval [ms] | 103 |
| Z-step [nm] | 50.0 |
| Camera offset [electron counts] | 91.0 |
| Read noise [electron counts] | 9.84 |
| **Find Maxima** | |
| Pre-Filter | None |
| Noise tolerance | 80 |
| **Fit Parameters** | |
| Dimensions | 1 |
| Filter | Simplex-MLE |
| Max Iterations | 500 |
| Box size [pixel] | 12.0 |
| Fix width | Not selected |
| **Filter Data** | Nothing selected |
| **Positions** | Only imaged at one position |
| **Skip Channels** | Not selected |

**Supplementary Table 10 |** Fitting parameters used in µManager's[16] 'Localization Microscopy' plug-in.
